# Utilizing gene co-expression networks with the rat kidney TXG-MAPr tool to enhance safety assessment, biomarker identification and human translation

**DOI:** 10.1101/2023.11.03.565444

**Authors:** Steven J. Kunnen, Giulia Callegaro, Jeffrey J. Sutherland, Panuwat Trairatphisan, Hugo W. van Kessel, Lukas S. Wijaya, Git Chung, Keith Pye, Keith M. Goldstein, Claire R. Teague, Ciaran P. Fisher, Julio Saez-Rodriguez, Colin Brown, Susan A. Elmore, Kathleen M. Heinz-Taheny, James L. Stevens, Bob van de Water

## Abstract

Toxicogenomic data represent a valuable source of biological information at molecular and cellular level to understand unanticipated organ toxicities. Weighted gene co-expression networks analysis can reduce the complexity of gene-level transcriptomic data to a set of biological response-networks useful for providing insights into mechanisms of drug-induced adverse outcomes. In this study, we have built co-regulated gene networks (modules) from the TG-GATEs and DrugMatrix rat kidney datasets consisting of time- and dose-response data for 180 compounds, including nephrotoxicants. Data from the 347 modules were incorporated into the rat kidney TXG-MAPr web tool, a user-friendly interface that enables visualization and analysis of module perturbations, quantified by a module eigengene score (EGS) for each treatment condition. Several modules annotated for cellular stress, renal injury and inflammation were statistically associated with concurrent renal pathologies, including modules that contain both well-known and novel renal biomarker genes. In addition, many rat kidney modules contain well annotated, robust gene networks that are preserved across transcriptome datasets, suggesting that these biological networks translate to other (drug-induced) kidney injury cases. Moreover, preservation analysis of human kidney transcriptomic data provided a quantitative metric to assess the likelihood that rat kidney modules, and the associated biological interpretation, translate from non-clinical species to human. In conclusion, the rat kidney TXG-MAPr enables uploading and analysis of kidney gene expression data in the context of rat kidney co-expression networks, which could identify possible safety liabilities and/or mechanisms that can lead to adversity for chemical or drug candidates.

**Translational Statement:** Gene co-expression networks (modules) were generated using rat kidney toxicogenomics data, which reduced data complexity and retained quantitative mechanisms to enhance safety assessment. Several stress, injury and inflammation modules were statistically associated with renal pathologies, useful for biomarker identification. Moreover, many rat kidney modules contained well-annotated, robust gene-networks that were preserved in human patients transcriptome data after renal transplantation, suggesting that these biological networks translate to human relevant kidney-injury. So, the rat kidney TXG-MAPr tool enables transcriptome analysis in the context of kidney co-expression networks, which could identify chemical-induced safety liabilities and/or mechanisms leading to adversity, relevant for human risk-assessment.

## Introduction

Kidneys play a pivotal role in drug disposition and excretion, making them an important target organ for adverse reactions. Chemical- or drug-induced kidney injury (DIKI) is often observed in (proximal) tubular epithelial cells, but other nephron segments like the glomerulus may also be targeted ^1^. Initial injury can lead to subsequent cell death followed by repair and regeneration or, if injury is severe, to loss of nephrons and progression to renal failure ^2^. The severity of the initial insult, the degree of cellular injury, and the potential for repair in these regions are determined by local extracellular drug concentration (blood and glomerular filtrate) and intracellular accumulation of drug driven by cellular transport/uptake. The balance between uptake and efflux transport, as well as drug metabolism dictate the effective exposure at cellular targets that yield on- or off-target pharmacology and toxicity^3^. Activation of cellular stress response pathways, including oxidative stress, DNA-damage and unfolded protein response (UPR), are early responses to injury and are critical in the adaptation versus progression of kidney injury, although the mechanistic understanding is still incomplete ^4,5^.

Nonclinical safety studies are designed to characterize target organ toxicity that can be monitored during clinical testing of drug candidates. Histopathology data is the gold standard for pre-clinical drug safety assessment, but only informs the occurrence of injury and not the mechanism ^6^. Association of drug-induced transcriptomic changes with histopathology findings may provide key mechanistic insights into biological responses leading to adversities. However, despite two decades of application, transcriptomic analysis has not achieved wide-spread application in drug safety assessment, perhaps in part due to difficulties in achieving meaningful qualitative and quantitative interpretation and useful safety predictions from high content transcriptomic data ^7,8^. Among the reasons for slow uptake is the difficulty in translating gene-level data into interpretable biological responses and the penalties incurred due to false discovery rates ^8,9^. However, recent publications demonstrate that co-regulated gene network approaches can organize high dimensional toxicogenomic data into a smaller set of biological response networks, which has been applied to uncover novel mechanisms underlying drug-induced liver toxicities ^10–13^.

In this study, we aimed to develop co-regulated gene networks to study the mechanism of kidney injury using toxicogenomic data. We applied weighted gene co-expression network analysis (WGCNA) to identify rat kidney specific co-regulated gene networks (modules) using the publicly available TG-GATEs (TG) and DrugMatrix (DM) toxicogenomic dataset from rat kidney ^14–17^. As done previously for primary human hepatocytes and rat liver ^10,11^, we have developed an R-shiny based interactive webtool, the rat kidney TXG-MAPr (https://txg-mapr.eu/login/). WGCNA gene modules could be quantitatively visualized by module eigengene scores (EGS) and reduced to meaningful biological responses relevant to understanding mechanisms of toxicity. Specific module responses were statistically associated with pathology phenotypes (clinical chemistry and histopathology), thereby linking cellular mechanisms to renal injury. Several of these modules linked to kidney injury are preserved across datasets and species, including kidney transcriptomic data from human patients ^18^. Our results suggest that gene co-regulated modules in the rat kidney TXG-MAPr can be applied to other toxicogenomic datasets for chemical and drug safety assessment and to identify modules and associated biological responses that are relevant in the context of human translational safety assessments.

## Methods

### Sample collection and gene expression analysis

Toxicogenomic data from the TG-GATEs (TG) rat kidney repository (Affymetrix Rat Genome 230 2.0 array) were downloaded ^16^ and jointly normalized using the Robust Multi-array Average (RMA) method within the *affy* package using R ^19,20^. The BrainArray chip description file (CDF) version 19 (http://brainarray.mbni.med.umich.edu/Brainarray/Database/CustomCDF/CDF_download.asp, Rat230_2 array version 19) was used to map Affymetrix probesets to NCBI gene Entrez_IDs ^21^, which resulted in 13877 unique probe sets, each mapped to a single gene. The TG kidney repository contains 975 rat treatments, where each treatment is defined as a combination of compound, time and concentration. Vehicle treated control samples are available for each individual time point (time-matched controls). The *limma* R package was used to calculate log2 fold-change (log2 FC) values, by building a linear model fit and computing differential expression by empirical Bayes moderation for each treatment condition ^22^. Toxicogenomic data from the DrugMatrix (DM) rat kidney repository and a renal ischemia reperfusion injury (IRI) study were downloaded from Gene Expression Omnibus (GEO; https://www.ncbi.nlm.nih.gov/geo/; <u>GSE57811</u> and <u>GSE58438</u>) and analysed following the same procedure as the TG data ^17,23^. For the DM data the time-matched vehicle control samples were indicated per treatment condition. The IRI study had one group of naïve treated rat controls (n=5), which was used for differential gene expression at the different timepoints after ischemia reperfusion. Human Affymetrix transcriptomics data (Human Genome U133 Plus 2.0 Array) from human patients after kidney transplantation were collected from GEO (<u>GSE36059</u>) ^18,24^, annotated using BrainArray CDF version 25 (HGU133Plus2) and normalized using RMA in the *affy* R package. One CEL file sample was removed because it was deprecated. Differential gene expression of each patient kidney transplant sample was contrasted with the eight nephrectomy controls samples using *limma* package.

For isolation of primary rat proximal tubular cells (PTC) 3 male Sprague Dawley rats (10 weeks old) were sacrificed and kidneys were harvested and decapsulated. The rats were supplied by Charles River, UK, and were housed at Newcastle University before being euthanized by cervical dislocation. Ethical approval for the isolation and culture of primary proximal tubules cells from rats was obtained from the Animal Welfare and Ethical Review Body at Newcastle University and all procedures were carried out in accordance with Animals (Scientific Procedures) Act 1986. Newcastle University holds OLAW Animal Welfare Assurance certification for animal welfare. At least 1 mg of whole kidney biopsy sample was soaked in 1 mL of RNAlater. Cortical slices were taken through the kidney samples from each animal and rat PTCs were isolated from the minced cortical slices as previously described ^25,26^. Approximately 1 million freshly isolated rat PTCs were soaked in 1 mL of RNAlater serving as the day 0 sample. The remaining PTCs were seeded onto 12-well Transwell inserts at density of 200,000 cells per insert. Medium was changed on day 1, 3 and 5 after seeding. PTCs were harvested day 1, 2, 5 and 7 post-seeding. On the day of RNA sampling, media was gently removed from the wells, membranes were cut out and put into 10 mL of RNAlater (Thermo Fisher) per timepoint. Samples were stored at 4°C overnight, then at −20°C. RNA was isolated from rat kidney samples and PTC cultures using Trizol + Direct-zol columns according to the referenced protocol (Zymo Research). For PTC cultures, RNA was isolated from Transwell filters (multiple filters were pooled per timepoint) and concentrated over RNA Clean & Concentrate (RCC) columns (Zymo Research). RNA quality was assessed by Agilent Bioanalyzer and all samples had a RIN > 6.3. For microarray analysis, 1 μg of total RNA was used for preparation of biotin-labeled cRNA. Samples were hybridized to Affymetrix Clariom-D rat arrays for transcriptome analysis and scanned on an Affymetrix GeneChip Scanner. Raw CEL files were annotated using BrainArray CDF version 25 (ClariomDRat) and normalized using RMA in the *affy* R package. Differential gene expression for the PTC cultures was contrasted with the whole kidney samples using the *limma* package to investigate the transcriptional response of cells taken into culture. Raw and processed data of the PTC cultures is deposited in EBML-EBI ArrayExpress database (<u>E-MTAB-13633</u>).

### Rat kidney gene co-expression network analysis

Kidney specific co-regulated gene networks (modules) were obtained from the TG and DM rat kidney log2 FC data matrix ^16,17^, using the *WGCNA* package in R ^15^. The rat kidney TG data contains transcriptomic data of single (3, 6, 9 and 24 hours) and daily repeated (4, 8, 15 and 29 days) exposures for 41 (drug) compounds at various dose levels (*i.e.,* 975 compound, dose and time combinations (conditions) in total). The rat kidney DM data contains transcriptomics data from daily repeated exposures with 139 (drug) compounds at 1 or 2 dose levels and for up to 14 days (*i.e.*, 360 treatment conditions). We applied unsigned WGCNA to the log2FC matrix of all TG and DM data to group co-induced and co-repressed genes together. The optimal soft power threshold of 8 was selected based on the standard power-law plotting within the *WGCNA* package and by maximizing the difference in basal gene expression of genes showing co-expression (module genes) versus genes not showing co-expression (excluded genes) based on a t-test with the assumption that low expressed genes are not correlating. From the 13877 unique genes on the Rat230_2 array, we obtained 399 co-expression modules containing 11,244 genes, while 2633 excluded genes did not meet the co-expression criteria. Modules were merged when having a pair-wise Pearson correlation ≥ 0.8 of the module eigengene scores (EGS) based on all treatments, which resulted in a final list of 347 merged modules (**Table S1**). Merged modules contain the suffix ‘m’, *i.e.,* module rat KIDNEY:2m (*i.e.,* rKID:2m) was merged from child modules 2 and 11. For each treatment condition a module EGS was computed as the first principal component of variation for all genes after Z-scoring the log2 FC matrix (**Table S2**), as described previously ^10,12^. This module EGS represents a quantitative measure of activation or repression of all genes (based on log2 FC) in the module. An EGS > 2 or < −2 was considered as a large perturbation in expression of the underlying genes ^13^. The correlation between the log2 FC of each gene with the module EGS was also calculated (called the corEG = correlation eigengene) to estimate the intramodular connectivity of the genes and their network (note that genes can also have opposite (negative) weighting because of the unsigned co-expression). The gene with the highest absolute corEG within a module is the hub gene and is the most representative gene of the module. The circular rat kidney TXG-MAPr dendrogram was constructed using the *ape* package based on Ward’s hierarchical clustering of pair-wise Pearson correlation for every module across all treatment conditions (**Figure S1**). Each main branch is indicated by letters A-I, followed by subbranches (*i.e.,* A1aIαi) as indicated in **Figure S1 and Table S1**. To improve visualization of the dendrogram, some module edges and nodes were manually adjusted to separate module clusters. The module EGS are displayed on the TXG-MAPr dendrogram as circles, with the size and colour proportional to the amount of induction or repression for a given treatment condition (**Figure 1A**, red to blue colour scale, respectively). Compound or treatment correlation was calculated for all treatment conditions using pair-wise Pearson or Spearman correlation of the module EGS. Similarly, module correlation was calculated across all modules (347 x 347 matrix). Preservation of the TG and DM rat kidney WGCNA module structure with other datasets (TG rat kidney only, DM rat kidney only, TG rat liver and GEO human kidney) was performed using the *modulePreservation* function within the *WGCNA* package in R ^27^. Modules with a Z-summary > 2 were considered moderately preserved and Z-summary > 10 were highly preserved. The R-script for WGCNA module generation and preservation analysis can be found in an online GitHub repository linked to persistent identifier in Zenodo: https://doi.org/10.5281/zenodo.14926143.

**Figure 1.**
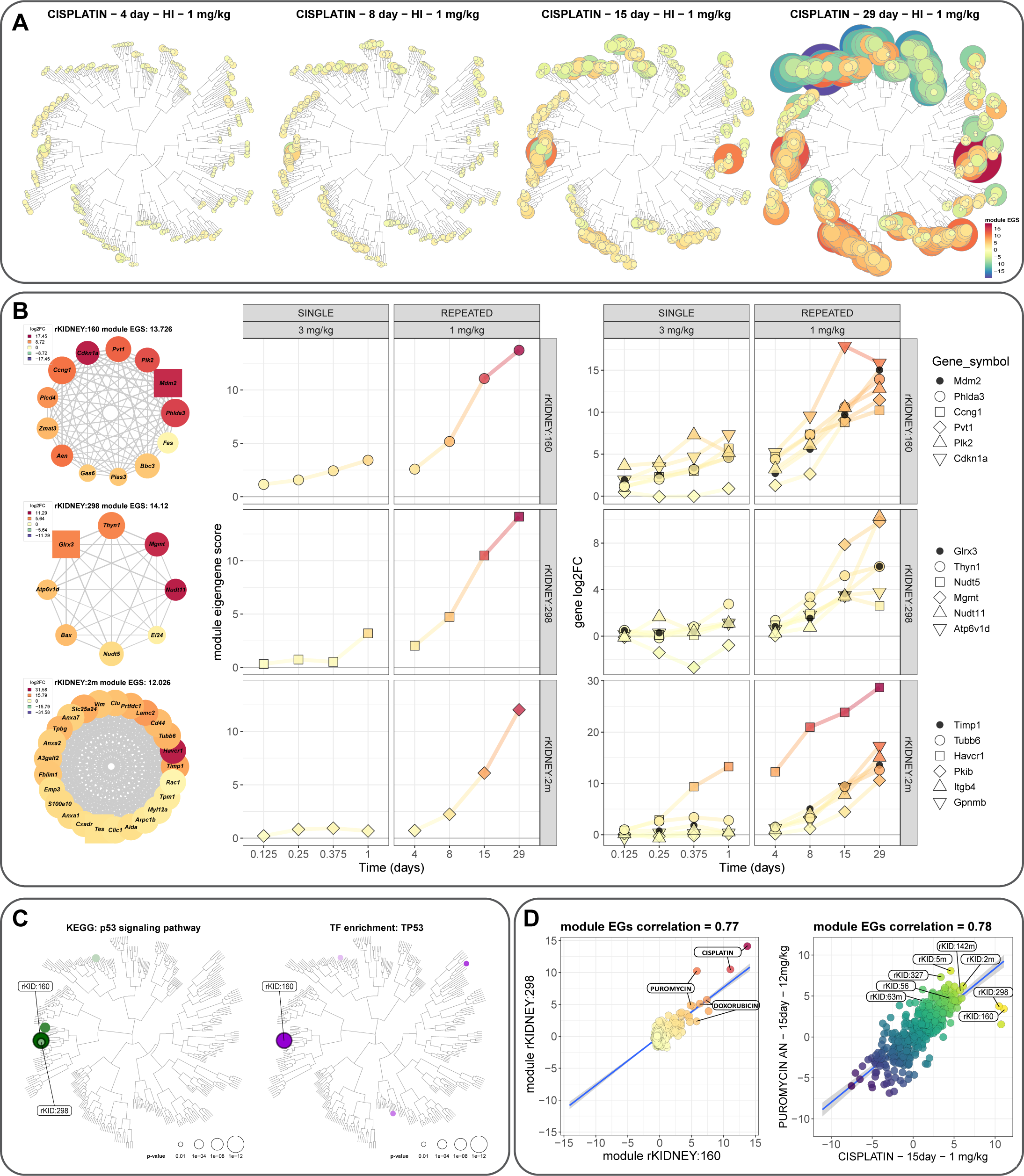
Overview of the rat kidney TXG-MAPr tool. (A) TXG-MAPr dendrograms showing WGCNA module eigengene scores (EGS) of daily repeated 1 mg/kg/day cisplatin time course exposure from 4 - 29 days. The size of the circles is proportional to the module EGS, and the red/orange colours indicate induction of the module, while blue/green indicates repression of the module. (B) Modules rKID:160, rKID:298 and rKID:2m (left) are strongly induced by high dose cisplatin exposure at 29 days, which is further displayed in cisplatin time response plots of the module EGS (middle) and the log2FC (right) of the most significant genes, including the most hub-like gene (black). (C) KEGG pathway and transcription factor (TF) enrichment (p-value) for p53 signalling or TP53 projected on the TXG-MAP dendrogram, with rKID:160 and rKID:298 highlighted. (D) Module EGS correlation plot of rKID:298 and rKID:160 for all treatment conditions (left) and correlation between 12 mg/kg/day puromycin aminonucleoside and 1 mg/kg/day cisplatin at 15 days for all modules (right).

### Functional enrichment analyses and annotation of modules

Modules were annotated by functional enrichment (GO-terms, pathways and transcription factors) using hypergeometric tests (**Tables S3-S5**) as described previously ^10^. Briefly, GO-term enrichment was performed with the *topGO* package in R using the algorithm = “classic” and statistic = “fisher” ^28^. Over Representation Analysis (ORA) was performed on the gene members of each module using Consensus Pathway DB (cpdb version 34), including the following databases: BioCarta, EHMN, HumanCyc, INOH, KEGG, NetPath, Reactome, PharmGKB, PID, Signalink, SMPDB, Wikipathways, UniProt, InterPro ^29^. For both resources, we included enriched terms with a hypergeometric test p-value < 0.01 (**Table S3**). To identify regulatory transcription factors (TFs) of each module, a hypergeometric test was performed on module gene members using the function *phyper* within the *stats* package in R (**Table S4**). The gene set of TFs and their regulated genes (regulons) are derived from DoRothEA ^30^ with two sets of confidence levels: the “high confidence” level comprises categories A, B and C, while the “high coverage” level comprises categories A, B, C and D. The enriched TFs with p-value less than 0.01 were included in the study. In parallel, TFs’ activities were estimated as normalized enrichment scores using the *viper* function from the *viper* package ^31^ with two confidence sets of TF-regulon from DoRothEA as described. All parameter settings were assigned as in the original DoRothEA study ^30^. The most significant module enrichment terms (gene overrepresentation analysis) were reviewed for a common theme, which were summarized in a Key_annotation and annotation levels 1 through 3 (**Table S5**). Only modules with an enrichment p-value < 10^-4^ (0.0001) were reviewed to find a common annotation. Annotation level 1 describes the general cellular process or function (*e.g.,* metabolism, immune response, RNA/protein processing, stress response), level 2 describes the sub-process or function, and level 3 provides the most specific process or transcription factor involved. These 3 levels of annotation were summarized in the Key_annotation, which describes the function of the genes in the modules based on the most significant enrichment terms (**Table S5**).

### Module association with pathology

The TG database also provides clinical chemistry (biomarker data) and histopathology data for all treatment conditions. Histopathology data was reviewed by pathologists for 4 nephrotoxic compounds (cisplatin, carboplatin, cyclosporine A and allopurinol) and most calls were in agreement with the pathology score provided by TG. Only the most occurring pathologies were included for the statistical association, including necrosis, degeneration, regeneration, basophilic change (cellular staining indication a regenerative process), cellular infiltration (indicating an inflammatory/ immune response), fibrosis, vacuolization, dilatation (widening of the renal tubules), cysts and hyaline casts (protein aggregates in the tubular lumen). Overlapping pathology terms were combined (*e.g.*, different types of cellular infiltration or different regions of tubular injury), resulting in a final list of 19 unique histopathology’s (**Tables S6-S7**). Necrosis and degeneration were also considered as a combined term because these pathologies have overlapping biological meaning, as well as regeneration and basophilic change. For these overlapping pathologies, the maximum grade was used for each treatment condition. Histopathology severity was converted in a numerical scale (normal = 0, minimal = 1, slight = 2, moderate = 3, severe = 4) and the average score was calculated per treatment group of maximum 3 animals that were also used for the microarray analysis. Histopathology presence in a treatment group was indicated by a severity score >= 0.67 for all pathologies, but a higher threshold (>=1.33) was also applied for a few pathologies (**Table S7**). Clinical chemistry measurements were calculated as average percent change from the time-matched control group. The KW-BW ratio (kidney weight - body weight ratio) was calculated by dividing the total kidney weight by the body weight. The threshold for a significant increase in clinical measurements is defined in **Table S7**. The kidney toxicity phenotypes (histopathology or clinical chemistry) were used to evaluate statistical association between module EGS and the occurrence of kidney pathology, as described previously ^13^. Both concurrent and predictive module associations with pathology (*i.e.*, early module changes that are preceding the toxicity at a later timepoint) were calculated. Effect sizes (Cohen’s D) of each module were calculated by comparing mean module EGS in the presence (positives) or absence (negatives) of renal injury, with thresholds specified in **Table S7**, for each of the 29 toxicity phenotypes (**Table S8**).

Cohen’s D = (mean module EGS positives – mean module EGS negatives) / pooled SD module EGS

To control for overall gene expression, we applied logistic regression to account for differences in both module EGS and average absolute EGS (AvAbsEGS) as covariates in the analysis, as described previously^13^. Briefly, the *glm()* function of the *stats* package was applied with single linear models:

Module_single = glm(phenotype ∼ module EGS, family = “binomial”)

AvAbsEGS_single = glm(phenotype ∼ AvAbsEGS, family = “binomial”)

The Module_single model predicted the outcome of using only module EGS as variable and provided p_single (**Table S8**). In addition, the full linear model was calculated:

Module_Av_full = glm(phenotype ∼ AvAbsEGS + module EGS, family = “binomial”)

This determined the coefficients, which is the natural log odds ratio for observing a phenotype for a single-unit increase in module EGS. Finally, the significance of module EGS as an additional variable was tested, which provided the p-adj (**Table S8**):

add1(AvAbsEGS_single, scope = ∼ AvAbsEGS + module EGS, test = “Chisq”)

The p-adj value was converted to a signed log10 p-adj, by using the sign of the effect sizes. This signed log10 p-adj and the Cohen’s D effect size were used as measure/rank for module associations with toxicity phenotypes in several plots and tables.

### Rat kidney TXG-MAPr webtool

An interactive visualization of rat kidney WGCNA module and gene expression data has been implemented in the rat kidney TXG-MAPr (Toxicogenomics-MAPr) webtool using the *shiny* package in R to create a user-interface ^32^, as described previously ^10^. The rat kidney TXG-MAPr tool is available at https://txg-mapr.eu/login/. The tool allows evaluation of dose- and time-response data, compound correlation plots and module association with pathology, which is also available in tabulated format. New or external gene expression data (log2 FC) can be uploaded in the TXG-MAPr tool, as described previously ^10^. This enables calculation of new EGS for each module from external dataset based on the gene log2 FC and weighted by the gene corEG. The new module EGS will be overlaid onto the rat kidney TXG-MAPr dendrogram and will be fully integrated into the web application for that session. Uploaded data will be removed when the session is closed.

### Quantification and statistical analysis

All figures were plotted in R version 4.0.0 ^19^or higher using mainly *ggplot2, pheatmap, shiny, igraph* and *ape* packages. Statistical analyses were carried out using functions from R packages for differential gene expression analysis (*affy*, *limma*), correlation analysis and logistic regression (*stats*), enrichment analysis (*topGO*, *stats*), gene co-expression and preservation analysis (*WGCNA*).

## Key resources table

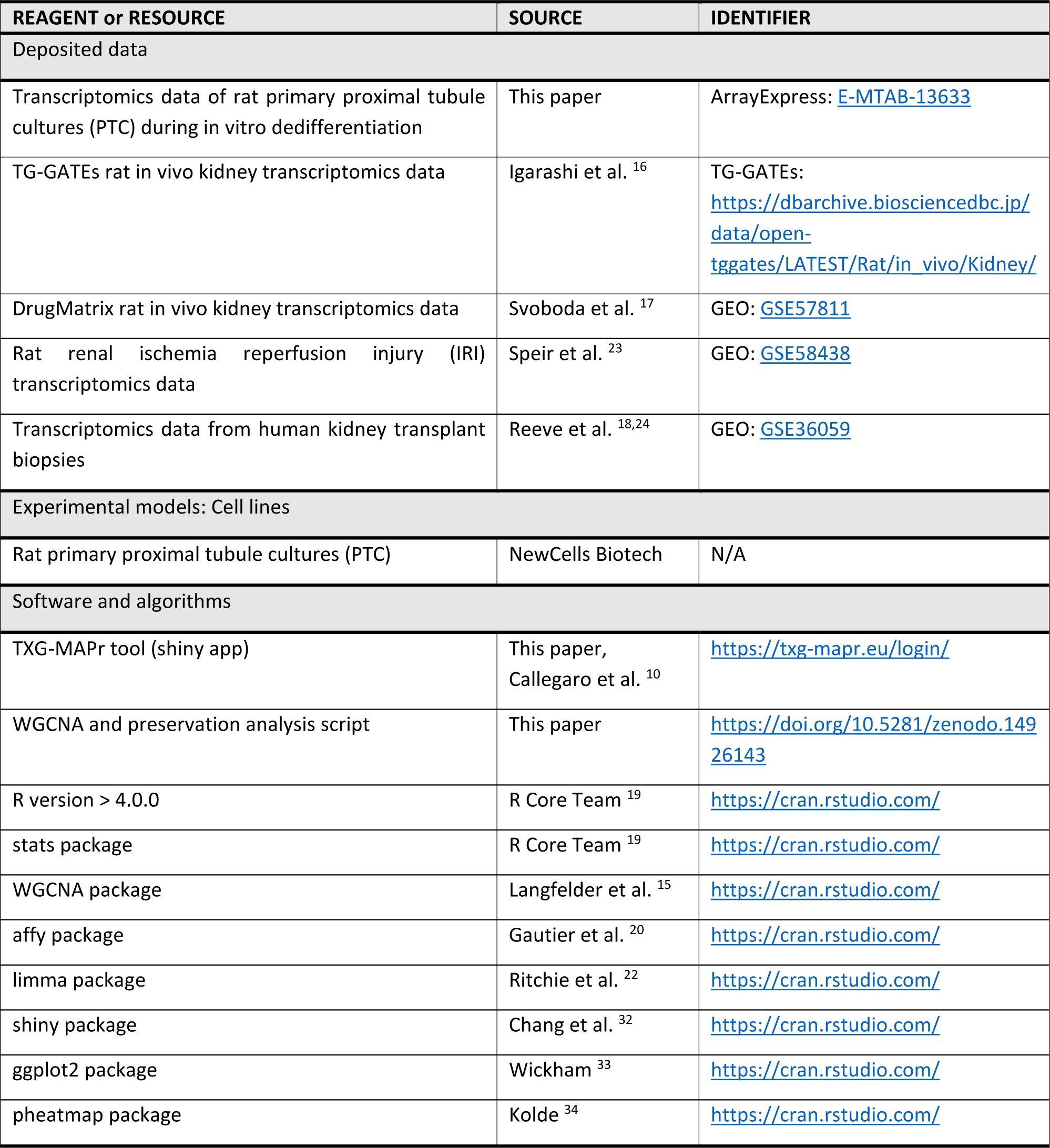

| REAGENT or RESOURCE | SOURCE | IDENTIFIER |
| --- | --- | --- |
| Deposited data |  |  |
| Transcriptomics data of rat primary proximal tubule cultures (PTC) during in vitro dedifferentiation | This paper | ArrayExpress: E-MTAB-13633 |
| TG-GATEs rat in vivo kidney transcriptomics data | Igarashi et al. <sup>16</sup> | TG-GATEs: https://dbarchive.biosciencedbc.jp/data/open-tggates/LATEST/Rat/in_vivo/Kidney/ |
| DrugMatrix rat in vivo kidney transcriptomics data | Svoboda et al. <sup>17</sup> | GEO: GSE57811 |
| Rat renal ischemia reperfusion injury (IRI) transcriptomics data | Speir et al. <sup>23</sup> | GEO: GSE58438 |
| Transcriptomics data from human kidney transplant biopsies | Reeve et al. <sup>18,24</sup> | GEO: GSE36059 |
| Experimental models: Cell lines |  |  |
| Rat primary proximal tubule cultures (PTC) | NewCells Biotech | N/A |
| Software and algorithms |  |  |
| TXG-MAPr tool (shiny app) | This paper, Callegaro et al. <sup>10</sup> | https://txg-mapr.eu/login/ |
| WGCNA and preservation analysis script | This paper | https://doi.org/10.5281/zenodo.14926143 |
| R version > 4.0.0 | R Core Team <sup>19</sup> | https://cran.rstudio.com/ |
| stats package | R Core Team <sup>19</sup> | https://cran.rstudio.com/ |
| WGCNA package | Langfelder et al. <sup>15</sup> | https://cran.rstudio.com/ |
| affy package | Gautier et al. <sup>20</sup> | https://cran.rstudio.com/ |
| limma package | Ritchie et al. <sup>22</sup> | https://cran.rstudio.com/ |
| shiny package | Chang et al. <sup>32</sup> | https://cran.rstudio.com/ |
| ggplot2 package | Wickham <sup>33</sup> | https://cran.rstudio.com/ |
| pheatmap package | Kolde <sup>34</sup> | https://cran.rstudio.com/ |

## Results

### Rat kidney TXG-MAPr tool to identify biological mechanisms of kidney injury

Gene co-expression networks (347 modules) were built from the TG and DM rat kidney datasets using WGCNA (**Table S1**). Gene expression changes were scored at the module level for each treatment condition by the eigengene score (EGS), see **Table S2** for all treatment conditions. The EGS indicates the induction or repression of the entire module based on the weighted log2 FC expression of its component genes. Modules are arranged in a TXG-MAPr dendrogram, based on the hierarchical clustering across all treatments (**Figure S1)**, and the size and colour of each circle is proportional to the amount of induction or repression for a given treatment condition (**Figure 1A**, red to blue colour scale, respectively). The rat kidney module and gene expression data were incorporated into a R-shiny framework (TXG-MAPr: https://txg-mapr.eu/login/) for interactive visualization and analysis, as previously described ^10^. Functional annotation of the modules was assessed using overrepresentation analysis (GO-term, pathway and transcription factor) to provide cellular and mechanistic context of gene members in each module (**Tables S3-S5**). Not all modules showed strong enrichment for GO and pathway terms, but others were highly enriched for terms relevant to kidney injury and regeneration, including rKID:3m (metabolism, e-45), rKID:5m (cell cycle, e-67), rKID:6m (ribosome biogenesis, e-42), rKID:7m (immune system, e-47) and rKID:10m (extracellular matrix, e-34). From this functional enrichment analysis, a specific key annotation term was derived for 170 modules (**Table S5**), *i.e.,* when annotation terms were highly significant (p < 0.0001) and show a similar theme or could be linked to the enriched transcription factor, which was included in further analyses (see below).

To illustrate the utility of the rat kidney TXG-MAPr tool as an approach to extract mechanistic information from gene expression data, we selected compounds in the TG dataset for which various pathologies were noted in the single and repeated dose studies (**Table S6**). Repeated daily dosing for up to 29 days with the genotoxic chemotherapeutic drug cisplatin (CSP) caused progressive perturbation of gene expression in rat kidneys, represented by the module induction or repression visualized on the TXG-MAPr dendrograms (**Figures 1A and S2**). Histology findings were also progressive over time, including tubular necrosis, dilatation and regeneration, and were accompanied by increases in blood urea nitrogen (BUN) and serum creatinine (CRE) (**Table S6**). We looked for activation of modules associated with the DNA-damage response, a primary mechanism of cisplatin inducing kidney injury ^35^. Rat kidney module 160 (*i.e.,* rKID:160) was among the top induced modules after cisplatin treatment at various timepoints and dose levels (**Table S2**) and contained many well-known p53-target genes (*Mdm2*, *Cdkn1a*, *Phlda3*, *Ccng1*, *Plk2, Aen*, *Zmat3*, *Fas*, *Pias3*, *Gas6*) involved in the DNA-damage response (**Figure 1B, top**), and had the highest transcription factor (TF) enrichment score for Tp53 (p-value of 2e-10) among all modules (**Figure 1C, Table S4**). Induction of module rKID:160 and its component genes was already evident early and increased over time (**Figure 1B, top**). Another module, rKID:298, also enriched for terms consistent with p53 signalling **(Table S3)** and contained genes involved in apoptosis (*Bax*, *Ei24, Thyn1*) and DNA repair (*Mgmt*), showed a similar pattern of induction compared to rKID:160 (**Figure 1B, middle**). Although both rKID:160 and rKID:298 are enriched for p53 signalling terms, only rKID:160 had strong TF enrichment for Tp53 (**Figure 1C, Table S4**). Nonetheless, module rKID:160 showed high correlation (Pearson R) with rKID:298 across all TG treatments (**Figure 1D, left**), indicating that both modules are strongly induced by the same treatment conditions (*e.g.,* especially genotoxic compounds cisplatin and doxorubicin), as expected given their proximity in the dendrogram and involvement in similar processes.

To determine if the modular gene expression responses also reflected the progressive injury noted with cisplatin treatment, we also investigated module membership for injury processes and kidney specific biomarkers. Module rKID:2m, contained well-known kidney biomarker genes, including kidney injury molecule-1 (Kim-1/*Havcr1)*, and was strongly associated with toxicity (discussed in the next sections). After only modest early induction, the rKID:2m EGS, and log2FC for component genes, increased markedly after day 4 (**Figure 1B, bottom**). Other modules that annotated for biological processes important for kidney injury and regeneration also showed dose and time dependent induction with cisplatin and other nephrotoxicants allopurinol (APL) and puromycin aminonucleoside (PAN) (**Figure S3**). Several modules involved in extracellular matrix (rKID:10m, 31m), immune response (rKID:56 and 63m), cytoskeleton (rKID:42m, 142m), cell cycle (rKID:5m) and transcription (rKID:29) were induced after 29 days cisplatin exposure and were also observed with other nephrotoxic compounds, including APL and PAN (**Figure S3**), both of which caused substantial kidney injury and regeneration (**Table S6**). Notably, the high module EGS correlation (Pearson R), between 15-days repeated treatment with 1 mg/kg/day cisplatin and 12 mg/kg/day puromycin aminonucleoside suggested that these responses reflect general kidney responses to injury, while also reflecting distinct mechanisms (**Figure 1D, right**). For example, the inflammatory (rKID:56, 63m and 327), cytoskeleton/ injury (rKID:2m and 142m) and cell cycle (rKID:5m) responses are comparable between CSP and PAN at day 15, but the p53 responses (rKID:160 and 298) are more pronounced with cisplatin, because of the genotoxic mechanism of action.

Taken together, the TXG-MAPr tool demonstrated that the co-expression modules capture meaningful biological responses after nephrotoxic insults, where early events, such as activation of p53 by cisplatin, are followed by later events associated with cellular and tissue injury, including modules containing inducible biomarker genes.

### Modules associate biological response networks with kidney pathology

Having demonstrated that modular gene expression captures valuable information on mechanism associated with kidney injury, we wanted to investigate the statistical association of module changes, and the associated biological responses, with kidney injury (histopathology or clinical chemistry, see **Table S6 and S7** for frequency and occurrence for each treatment). Therefore, for each relevant kidney toxicity phenotype (**Table S7**), we measured effect sizes (Cohen’s D) of module EGS for treatments resulting in the presence (positives) or absence (negatives) of renal injury (**Table S8**). To control for the association of injury with overall gene expression, we also applied logistic regression to account for differences in module EGS while controlling for average absolute EGS (AvAbsEGS) as covariates in the analysis, as described previously ^13^. Estimation of the odds of toxicity was quantified by the (adjusted) p-value from the logistic regression, which was signed (signed log10 p-adjust) for positive (induction) or negative (repression) module responses associated with toxicity. Both concurrent and predictive (*i.e.,* early module changes that precede emergence of toxicity at a later timepoint) module associations with pathology were calculated, although concurrent associations showed higher statistical significance as expected (**Table S8**).

The effect sizes of many modules associated with concurrent toxicity phenotypes showed strong Pearson correlation, suggesting that several biological response-networks are associated with multiple toxicity phenotypes (**Figure 2A**). This is likely an effect of either the high transcriptional activity, measured as AvAbsEGS, concurrent with multiple pathologies, and/or activation of generalized tissue responses to injury. When we repeated the clustering using the signed p-adjust values, which accounts for the AvAbsEGS as a covariate in the logistic regression, there was somewhat better separation of the toxicity phenotypes into two distinct clusters (**Figure 2B**). Cluster 1 modules were more associated with tubular injury phenotypes (increases in tubular necrosis, degeneration, BUN and CRE), and cluster 2 with regeneration and inflammation phenotypes (increases in tubular regeneration, dilatation, cellular infiltration and fibrosis). The overlap and differences in module association could be appreciated when comparing the TXG-MAP dendrogram plots overlaid with the signed log10 p-value per module for the most prevalent renal injuries (**Figures 2C and S4**). Modules in branch (see branches in **Figure S1**) F1 and F2 (stress responses and cell cycle) were strongly associated with tubular injury phenotypes (red colours). In contrast, modules in branch A2 and B2 (immune response and extracellular matrix / adhesion) were strongly associated with regeneration and inflammation. Cellular stress, transcription and cytokine related modules in branch H1 showed strong association with all pathologies (red colours), as well as modules in branch D2a (RNA splicing). Modules in branch I (mitochondrion/respiration) showed negative association with all toxicity phenotypes, as well as some metabolism modules in branch H1 (blue colours). Other (non-tubular) injuries did not show strong association with transcriptional module changes, likely due to the low number of positive treatments in the TG data that cause these injuries or the limited transcriptional perturbation (**Figure S4**).

**Figure 2.**
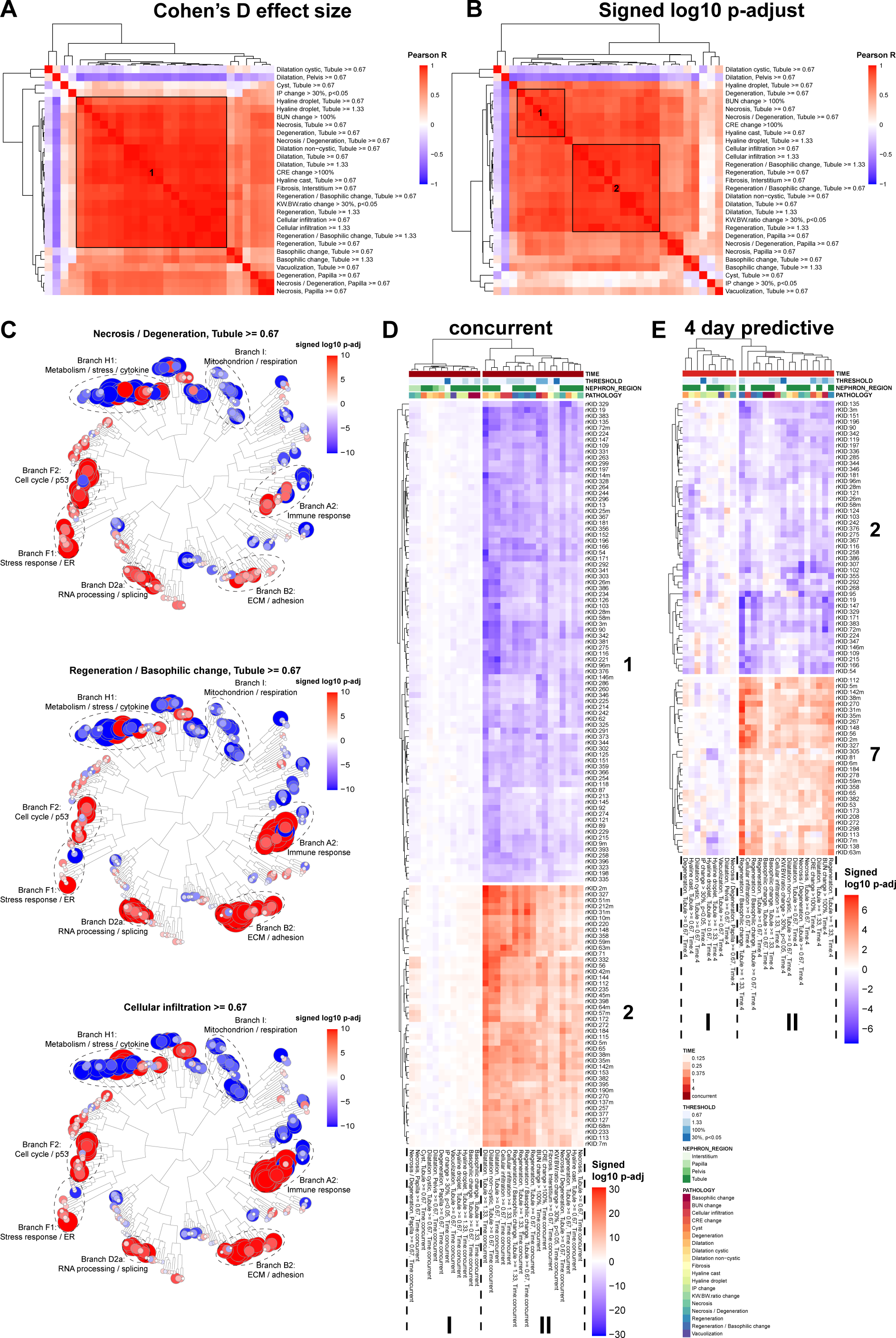
Module association with pathology. Strong Pearson correlation between different pathologies based on Cohen’s D effect size (A) and signed log10 adjusted p-value (B). This indicates that the same modules are associated with multiple pathologies, which could be the results of co-occurring pathologies (cluster 1 Figure A). Cluster 1 from figure A could be separated in 2 clusters (B), when controlling for module EGS using signed log10 adjusted p-values. Only the most occurring pathologies were taken along, with a pathology score ≥ 0.67 (and ≥ 1.33 for some pathologies). (C) TXG-MAPr dendrograms showing module association with different toxicity phenotypes based on the signed log10 adjusted p-value. Red colour indicates that the module EGS positively correlates with the selected pathology (module induced with pathology present), while blue means a negative correlation (module repressed with pathology present). The size of the circles is proportional to the -log10 adjusted p-value. For the branches with the strongest pathology associations, the general enrichment score is provided based on ORA of all genes in the branch. (D-E) Heatmap of signed log10 p-adjust of concurrent (D) and 4-day predictive (E) module associations with pathology. Red colour indicates that the module EGS positively correlates with the selected pathology, while blue means a negative correlation (*i.e.,* module is repressed when the pathology is present). Hierarchical clustering was applied on rows and columns to cluster both modules (rows) and toxicity phenotypes (columns) with similar association scores together. For concurrent associations, modules from clusters 1 and 2 were selected for strongest negative and positive correlation with toxicity, respectively. Module shows highest statistical association with concurrent toxicity phenotypes (see bottom pathology cluster II), which were selected to look at the mean effect sizes and p-values (see also **Table S9**). For predictive associations, the 4 days show the highest statistical significance, where modules from clusters 2 and 7 were selected for strongest negative and positive correlation with toxicity, respectively. Heatmaps are clustered by Euclidean distance, using the complete method from *pheatmap* package.

Hierarchical clustering of the modules for concurrent association with toxicity phenotypes revealed a subset of 44 strong positively (cluster 2) and 81 strong negatively (cluster 1) associated modules with injury and repair phenotypes (**Figures 2D, S5A**, **Table S9**), including serum biomarkers increases (BUN or serum creatinine), several types of tubular injury, regeneration, inflammation or fibrosis (pathology cluster II in **Figure 2D and S5A**, **Table S7**). Notably, module rKID:2m, which contained genes for known and inducible biomarkers of renal injury, is in cluster 2 (**Figure 2D**) and ranked highest, on average, for concurrent association with the most prevalent toxicities on both effect size and adjusted p-value (**Table S9**). Other positively associated and high ranked modules (cluster 2, **Figure 2D and S5A, Table S9**) were annotated for immune response (rKID:56, 63m, 71, 212m and 327), cytoskeleton (rKID:42m, 112, 142m), extracellular matrix organization (rKID:10m, 31m), proteasome (rKID:35m), cell cycle (rKID:5m) and transcription (rKID64m), all of which are processes associated with progressive renal injury and/or repair ^36,37^, and were induced by compounds like cisplatin, allopurinol and puromycin aminonucleoside which caused progressive renal injury (**Figure S3**). Negatively associated and high ranked modules (cluster 1, **Figure 2D and S5A, Table S9**) were annotated for metabolism (rKID:3m, 9m, 14m, 25m, 292), transport (rKID:19, 171, 329), AMPK signalling (rKID:383), mitochondrion (rKID:26m, 28m, 96m, 264) and renal or glomerular development (rKID:196) amongst others. Interestingly, the latter module rKID:196 showed various genes involved in glomerular development or podocyte function, including Nphs1, Podxl, Ptpro and Wt1^38,39^.

Transcriptional changes at day 4 showed on average the highest statistical power (effect size and log10 adjusted p-value) to predict renal injury on a later timepoint, followed by day 1 (**Figure S6**). Modules for predictive association with toxicity phenotypes at day 4 revealed a subset of 30 positive (cluster 7) and 46 negative (cluster 2) associations (**Figures 2E and S5B**, **Table S9**). Predictive association at day 1 revealed a subset of 44 positive (cluster 3) and 17 negative (cluster 1) associations (**Figure S5B**, **Table S9**). Several modules that showed strong positive predictive association with pathology were also strongly associated with concurrent pathology, including cytoskeleton (rKID:2m, 112, 142m) and immune response (rKID:7m, 56, 63m, 113, 212m, 327), while mitochondrial (rKID:26m, 28m and 96m) and metabolism (rKID:3m) showed negative association. Besides the overlap between modules that showed both concurrent and predictive association with pathology, there were also some unique modules in the predictive association selection. These include modules involved in ribosome biogenesis (rKID:6m, 8m, 40m, 130m), RNA processing (rKID:34m, 278, 387), protein processing (rKID:173, 208), immune response (rKID:16m, 138, 351), transcription (rKID:29), p53 pathway (rKID:160, 298), Nrf2 pathway (rKID:53), Atf4 pathway (rKID:74) and Atf6 pathway (rKID:81), which are early responses after cellular stress or injury.

Overall, this led to a subset of 169 modules (75 positive and 94 negative) that showed strongly perturbed transcriptional activity and could be correlated with induction of renal injury (**Table S9,** column “selected strong association”). Thus, module changes reflect biological responses that may be associated with higher likelihood of concurrent or future development of kidney adverse outcomes (*i.e.,* pathologies), while these modules may also contain potential biomarkers.

### Pathology associated modules facilitate biomarker discovery

In the previous section, we have described several modules that were most strongly associated with pathogenesis (**Table S9**), including rKID:2m, which contained well-known kidney injury biomarkers, including kidney injury molecule-1 (Kim-1/*Havcr1)*, lipocalin 2, (*Lcn2*/Ngal); osteopontin (*Spp1)*, clusterin (*Clu)* and tissue inhibitor of metalloproteinases (*Timp1*), all of which had high corEG (*i.e.,* hubness) (**Table 1**). Interestingly, the primary enrichment terms for rKID:2m included cell adhesion and actin cytoskeletal organization, suggesting a more complex set of biological responses is associated with activation and interpretation of the inducible biomarker genes in rKID:2m. Other known kidney injury biomarkers were in metabolism (rKID:3m), immune response (rKID:7m, 15, 56, 63m and 281) and extracellular matrix (rKID:10m) modules, and most were strongly associated with renal pathogenesis (**Tables 1, S9**).

**Table 1.** Kidney injury biomarkers in co-expression modules. Well-known and established kidney injury biomarkers are in cytoskeleton / injury (rKID:2m), metabolism (rKID:3m), immune response (rKID:7m, 15, 56, 63m and 281) and extracellular matrix (rKID:10m) modules.

| Gene | Biomarker name | Injury marker | Module | corEG | References |
| --- | --- | --- | --- | --- | --- |
| **Havcr1** | KIM-1 | Proximal Tubular | rKID:2m | 0.8156 | 40–46 |
| **Lcn2** | Lipocalin2, NGAL | Proximal / Distal Tubular | rKID:2m | 0.7698 | 42–46 |
| **Clu** | Clusterin | Proximal / Distal Tubular | rKID:2m | 0.8282 | 41,45,46 |
| **Spp1** | Osteopontin, OPN | Inflammation / Distal Tubular | rKID:2m | 0.7815 | 42,46 |
| **Timp1** | Timp-1 | Proximal Tubular | rKID:2m | 0.8948 | 47,48 |
| **Umod** | Uromodulin | Loop of Henle | rKID:3m | 0.7673 | 42,44 |
| **Tff3** | Trefoil Factor 3 | Proximal Tubular | rKID:3m | 0.5565 | 41 |
| **Cst3** | Cystatin C | Glomerular / Proximal Tubular | rKID:7m | 0.3903 | 41,43–46 |
| **B2m** | β2-microglobulin | Glomerular / Proximal Tubular | rKID:7m | 0.6849 | 41,45 |
| **Timp2** | Timp-2 | Proximal Tubular | rKID:10m | 0.7746 | 42,44 |
| **Igfbp7** | IGFBP-7, IBP-7 | Proximal Tubular | rKID:10m | 0.5295 | 42,44 |
| **Ambp** | α1-microglobulin | Glomerular / Proximal Tubular | rKID:15 | 0.7985 | 46 |
| **Alb** | Albumin | Glomerular / Proximal Tubular | rKID:15 | 0.7157 | 41,43,45,46 |
| **Fabp1** | L-FABP | Glomerular / Proximal Tubular | rKID:15 | 0.8531 | 42,44,45 |
| **Tnfrsf1a** | TNFR1 | Inflammation | rKID:56 | 0.9015 | 49 |
| **Tnfrsf1b** | TNFR2 | Inflammation | rKID:63m | 0.6241 | 49 |
| **Il18** | Interleukin-18 | Inflammation | rKID:63m | 0.7694 | 42–46 |
| **Ccl2** | MCP-1 | Inflammation | rKID:281 | 0.7258 | 42,44 |

To discover novel inducible kidney injury biomarker genes, we investigated the modules with strong pathology association and clear biological annotation. The top 10 modules showing the strongest positive association with each of the eight most severe kidney pathologies were selected, which resulted in a set of 29 unique modules (**Tables 2, S9**). The selected modules showed very good overlap with modules in cluster 2 of **Figure 2D and S5A** (concurrent), as well as with cluster 6 and 7 in **Figure 2E and S5B** (4-day predictive). In addition to the known biomarker genes found in these modules (**Table 1**), many modules with strong pathology association contain hub(-like) genes of which literature evidence indicates that their respective proteins could be potential kidney injury or disease biomarkers (**Table 2**). Modules with potential biomarkers are rKID:7m (*Cst3* = Cystatin C*, B2m* = β2-microglobulin*, Cd68*), rKID:10 (*Cdh11, Smoc2*), rKID:31m (*Il24*), rKID:38m (*Tcp1*), rKID:56 (*Icam1, Tnfrsf1a*), rKID:63m (*C1qb, Il18, Tnfrsf1b*), rKID:71 (*Mmp7*), rKID:138 (*Cd3g*), rKID:142m (14-3-3 proteins, *i.e., Ywhah, Ywhag*), rKID:212m (*Cp* = ceruloplasmin, *C1s, C1r, Dcn*) and rKID:327 (fibrinogen, *i.e., Fga, Fgb, Fgg*). Many genes in these modules are strongly induced by nephrotoxic treatments, including the potential biomarker genes (**Figure 3A**). Gene expression responses of these modules mimicked the time-dependence of kidney pathology (**Figure 3B**), suggesting that these modules could be relevant for characterization of previously identified biomarkers and identification of novel biomarkers of kidney injury. In contrast, traditional serum biomarkers (BUN and CRE) increased much later (**Figure 3C**), when there is loss of renal function and hence these do not recapitulate the renal injury already present at earlier timepoints. These results suggest that induction of putative biomarker genes and their modules is strongly associated with renal pathology and capture the progression of kidney injury much better than serum biomarkers BUN or CRE.

**Figure 3.**
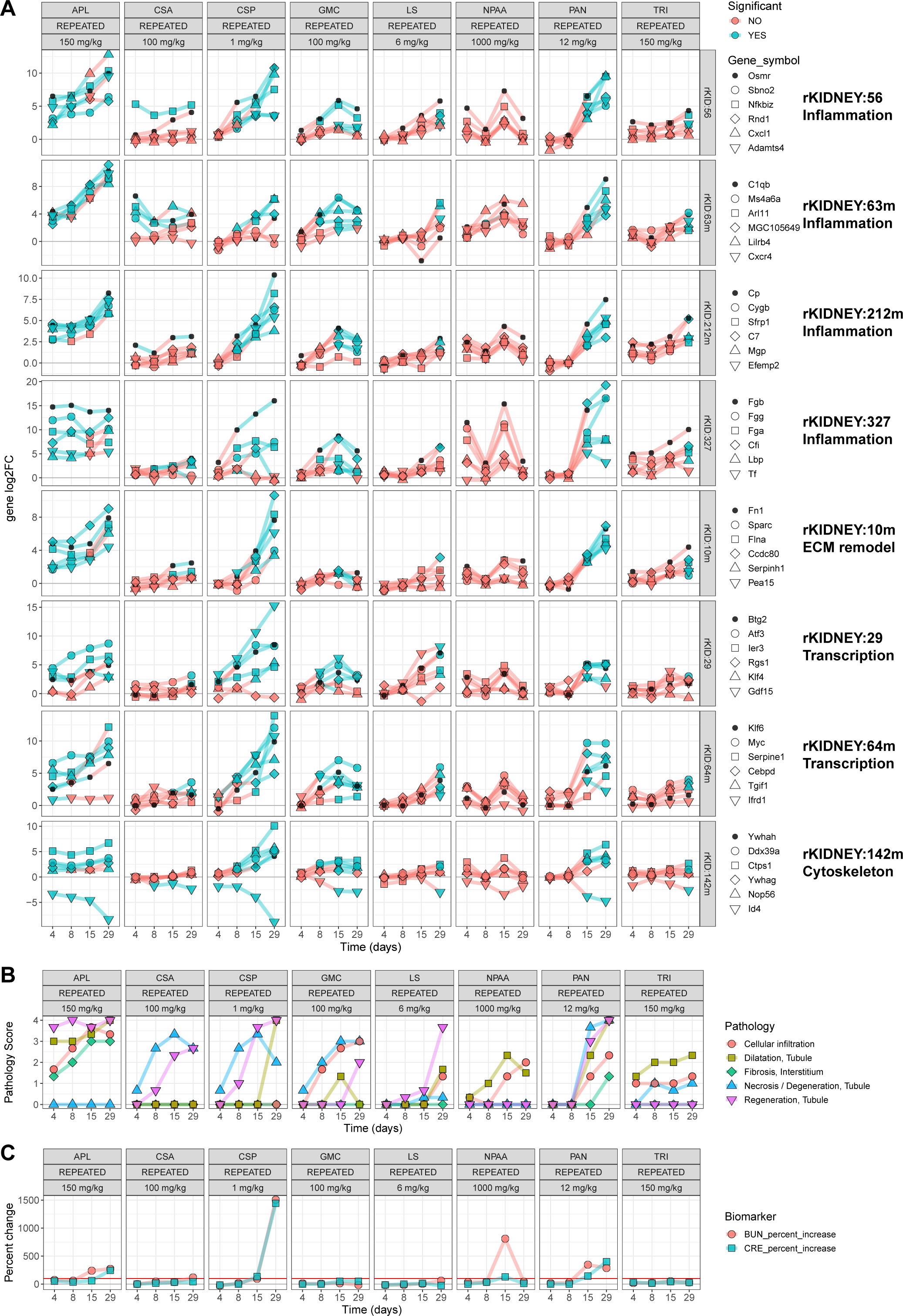
Gene expression responses during nephrotoxic treatments. (A) Expression of genes of selected modules with high pathology association is induced during nephrotoxic treatment conditions. (B) The most prevalent renal pathologies induced by nephrotoxic treatments. (C) Serum biomarker increases during nephrotoxic treatments versus controls (percentage increase, red line indicates 100% or 2-fold increase). APL = allopurinol, CSA = cyclosporine A, CSP = cisplatin, GMC = gentamicin, LS = lomustine, NPAA = phenylanthranilic acid, PAN = puromycin aminonucleoside, TRI = triamterene.

**Table 2.** Most strongly associated modules with renal pathology containing novel biomarkers. Top 10 ranked modules for positive association with concurrent kidney pathologies were selected for each phenotype. For each module, the hub gene and other highly connected genes are shown with a focus on identification of the corresponding proteins as putative biomarkers of renal injury (see references). The module annotations provide additional mechanistic context for these candidate biomarkers. The strongest pathology association, the rank and the signed log10 p-adjust value is provided for each module. Some of the modules are in the top 10 rank for more than one toxicity phenotype (column: “# Toxicity phenotypes”), including rKID:2m, 31m, 71 and 327.

| Module | Module annotation | Hub gene | Other hub genes | Branch | Strongest pathology association | # Toxicity phenotypes | Rank | Max signed log10 p-adjust | References (DOI) |
| --- | --- | --- | --- | --- | --- | --- | --- | --- | --- |
| rKID:31m | Extracellular matrix (ECM) | Il24 |  | A2allβ | Regeneration / Basophilic change, Tubule >= 0.67 | 3 | 1 | 27.68 | [10.1007/s00795-019-00239-0](https://doi.org/10.1007/s00795-019-00239-0) |
| rKID:212m | Immune response; complement | Cp | Ceruloplasmin (Cp), C1s, C1r, Dcn | B2all | Regeneration / Basophilic change, Tubule >= 0.67 | 1 | 2 | 23.94 | [10.1007/s10157-019-01734-5](https://doi.org/10.1007/s10157-019-01734-5)<br>[10.1681/ASN.2008111145](https://doi.org/10.1681/ASN.2008111145) |
| rKID:358 |  | Fkbp10 |  | B2all | Regeneration / Basophilic change, Tubule >= 0.67 | 2 | 3 | 22.49 |  |
| rKID:148 |  | Sept10 |  | D2a | Regeneration / Basophilic change, Tubule >= 0.67 | 1 | 5 | 19.54 |  |
| rKID:10m | Extracellular matrix (ECM) | Fbn1 | Cdh11, Smoc2 | B2all | Regeneration / Basophilic change, Tubule >= 0.67 | 1 | 8 | 18.68 | [10.1016/j.kint.2021.04.037](https://doi.org/10.1016/j.kint.2021.04.037) |
| rKID:184 | Other | Rexo2 |  | F2b | Necrosis / Degeneration, Tubule >= 0.67 | 1 | 2 | 18.83 |  |
| rKID:142m | Cytoskeleton | Ywhah | 14-3-3 protein (Ywhah, Ywhag) | H1bl | Necrosis / Degeneration, Tubule >= 0.67 | 2 | 5 | 17.09 | [10.1111/jcmm.13691](https://doi.org/10.1111/jcmm.13691) |
| rKID:38m | Protein processing; folding | Cct5 | Tcp1 | F1al | Necrosis / Degeneration, Tubule >= 0.67 | 2 | 7 | 15.80 | [10.1093/toxsci/kfz242](https://doi.org/10.1093/toxsci/kfz242) |
| rKID:35m | Proteasome | Eef1d |  | F2b | Necrosis / Degeneration, Tubule >= 0.67 | 1 | 10 | 15.51 |  |
| rKID:112 | Cytoskeleton | Arpc3 |  | F2al | Hyaline cast, Tubule >= 0.67 | 1 | 6 | 11.71 |  |
| rKID:5m | Cell cycle; S/M phase | Mad2l1 | Aurkb | F2b | Hyaline cast, Tubule >= 0.67 | 1 | 7 | 11.70 | [10.2147/IJGM.S336799](https://doi.org/10.2147/IJGM.S336799) |
| rKID:220 | Signalling | Htra3 |  | A2allβ | Fibrosis, Interstitium >= 0.67 | 1 | 5 | 9.33 |  |
| rKID:272 |  | Cdk4 |  | F2b | Fibrosis, Interstitium >= 0.67 | 1 | 10 | 8.75 |  |
| rKID:2m | Cytoskeleton; cell adhesion / injury biomarkers | Tes | Havcr1, Clu, Spp1, Lcn2, Timp1, Plaur | H1allαii | Dilatation, Tubule >= 0.67 | 4 | 2 | 25.13 | [10.1097/MNH.0000000000000275](https://doi.org/10.1097/MNH.0000000000000275) |
| rKID:51m |  | Trpv6 |  | A2allα | Dilatation, Tubule >= 0.67 | 3 | 3 | 25.06 |  |
| rKID:327 | Immune response; complement and coagulation | Fgb | Fibrinogen (Fga, Fgb, Fgg) | H1allαii | Dilatation, Tubule >= 0.67 | 4 | 4 | 24.50 | [10.1016/j.ajpath.2012.06.004](https://doi.org/10.1016/j.ajpath.2012.06.004) |
| rKID:235 |  | Tspan8 |  | H2al | Dilatation, Tubule >= 0.67 | 1 | 5 | 24.49 |  |
| rKID:144 |  | Pla1a |  | H1allαii | Dilatation, Tubule >= 0.67 | 2 | 6 | 24.25 |  |
| rKID:332 | Signaling; calcium | Map3k1 |  | H2all | Dilatation, Tubule >= 0.67 | 1 | 7 | 24.05 |  |
| rKID:56 | Immune response; TNF/NFκB signaling | Icam1 | Tnfrsf1a | H1blI | Dilatation, Tubule >= 0.67 | 2 | 9 | 22.68 | [10.1159/000068920](https://doi.org/10.1159/000068920)<br>[10.1097/MNH.0000000000000275](https://doi.org/10.1097/MNH.0000000000000275) |
| rKID:71 | Immune response | Mmp7 |  | A2allβ | Dilatation, Tubule >= 0.67 | 4 | 10 | 22.36 | [10.1681/ASN.2016030354](https://doi.org/10.1681/ASN.2016030354) |
| rKID:63m | Immune response | C1qb | Il18, Tnfrsf1b | A2b | Cellular infiltration >= 0.67 | 2 | 1 | 27.36 | [10.1111/petr.13885](https://doi.org/10.1111/petr.13885) |
| rKID:7m | Immune response; leukocyte activation | Ptpcr | B2m, Cd68, Coro1a, Cst3 | A2b | Cellular infiltration >= 0.67 | 1 | 4 | 21.23 | [10.1111/his.13768](https://doi.org/10.1111/his.13768) |
| rKID:138 | Immune response; T-cell receptor | Cd3g | Cd3 (Cd3d, Cd3e, Cd3g) | A2b | Cellular infiltration >= 0.67 | 1 | 5 | 21.23 | [10.1111/cei.12461](https://doi.org/10.1111/cei.12461) |
| rKID:233 |  | Tmsb4x |  | B2all | Cellular infiltration >= 0.67 | 2 | 8 | 17.86 |  |
| rKID:305 | Immune response; T-cell receptor | Zap70 | Cd247, Klrk1 | A2b | Cellular infiltration >= 0.67 | 1 | 9 | 17.81 |  |
| rKID:59m |  | Akip1 |  | B2all | Cellular infiltration >= 0.67 | 1 | 10 | 17.76 | [10.1097/MD.00000000000020742](https://doi.org/10.1097/MD.00000000000020742) |
| rKID:29 | Transcription | Btg2 | Atf3 | H1bl | BUN change > 100% | 1 | 9 | 15.51 | [10.21037/atm-21-5231](https://doi.org/10.21037/atm-21-5231) |
| rKID:64m | Transcription | Klf6 |  | H1bl | BUN change > 100% | 1 | 10 | 15.24 | [10.1080/0886022X.2020.1793353](https://doi.org/10.1080/0886022X.2020.1793353) |

### Module correlation analysis reveals both mechanistic and generalized tissue responses to kidney injury

Results already discussed allude to the fact that kidney injury and progression occur in distinct phases. Early molecular initiating events may be mechanistically distinct for different nephrotoxicants, but lead to a common event, such as death of proximal tubule epithelial cells, which results in a regeneration response. Therefore, we further explored how treatment correlation analysis, using module EGS, might reflect on early and later events in the progression of kidney injury. Besides cisplatin, other potentially nephrotoxic compounds show strong perturbations of gene expression and pathology at the highest tested dose levels and later timepoints (**Figures 3, S3, S7 and S8**). As exemplified earlier, there was a good correlation (R > 0.7) among module EGS after 15-days cisplatin and puromycin aminonucleoside treatment (**Figure 1D, right**), as well as after 8- and 29-days treatment (**Figure S9A**). Both drug treatments also induced comparable pathologies at these later timepoints, including tubular necrosis, dilatation and regeneration (**Figures 3B and S8B**, **Table S6**). At the 3-hour timepoint there was also a modest correlation (R = 0.55) between CSP and PAN, mainly driven by transcriptional stress (rKID:29 and 64m) responses (**Figure S9A**). However, from 6 hours through 4 days post treatment there was poor correlation of module EGS between CSP and PAN and little indication of pathology (**Figures S8B, S9A**). Induction of the p53 response by CSP is apparent at day 1 and later (rKID:160 and 298), while PAN induced modules associated with an early inflammatory response (rKID:15, 16m, 63m, 113, 351), the anticipated mechanisms to induce glomerular nephropathy ^50^.

Correlation analysis based on module EGS was further leveraged to investigate similarities between nephrotoxic treatment conditions given their overlap in activated and deactivated modules (**Figures 4, S8A and S9**). Treatment with nephrotoxic compounds at high dose levels and late time points showed strong correlation and clustering, when there was clearly pathology present in the kidneys (**Figure 4A cluster II, S9B**). This suggests that gene expression patterns are similar when there are co-occurring pathologies, likely an effect of upregulation of injury induced repair mechanisms of the tissue (*e.g.*, regeneration, dedifferentiation and inflammation responses), and downregulation of metabolism and mitochondrial function, that may be common after acute drug induced kidney injury ^51^. However, mechanistic information of compound exposures can still be extracted by focusing on outlier modules (rKID:160 and 298 in **Figure 1D**) or earlier timepoints when module activities are more diverse, see distinct compound specific clusters (**Figure 4A**, cluster I = APL, III = CSP, IV = PAN). The DM data also showed clustering of stress, immune, injury and regeneration responses with high module EGS during nephrotoxic exposures that clearly showed strong pathology association (**Figure S10, cluster 6 and 7, see row annotation**). Mitochondrial and metabolism modules that were negatively associated with pathology were indeed downregulated during these exposures with nephrotoxicants in the DM data (**Figure S10, cluster 1, 2 and 5**). Two estrogen receptor agonists primarily induced steroid hormone metabolism modules amongst others, showing their primary mechanism of action (**Figure S10, cluster 9**), while exposure with diethylstilbestrol showed clear activation of stress and injury response modules (cluster 6 and 7), which leaded to mild renal pathology on day 5 (**Figure S10B, see column annotation**).

**Figure 4.**
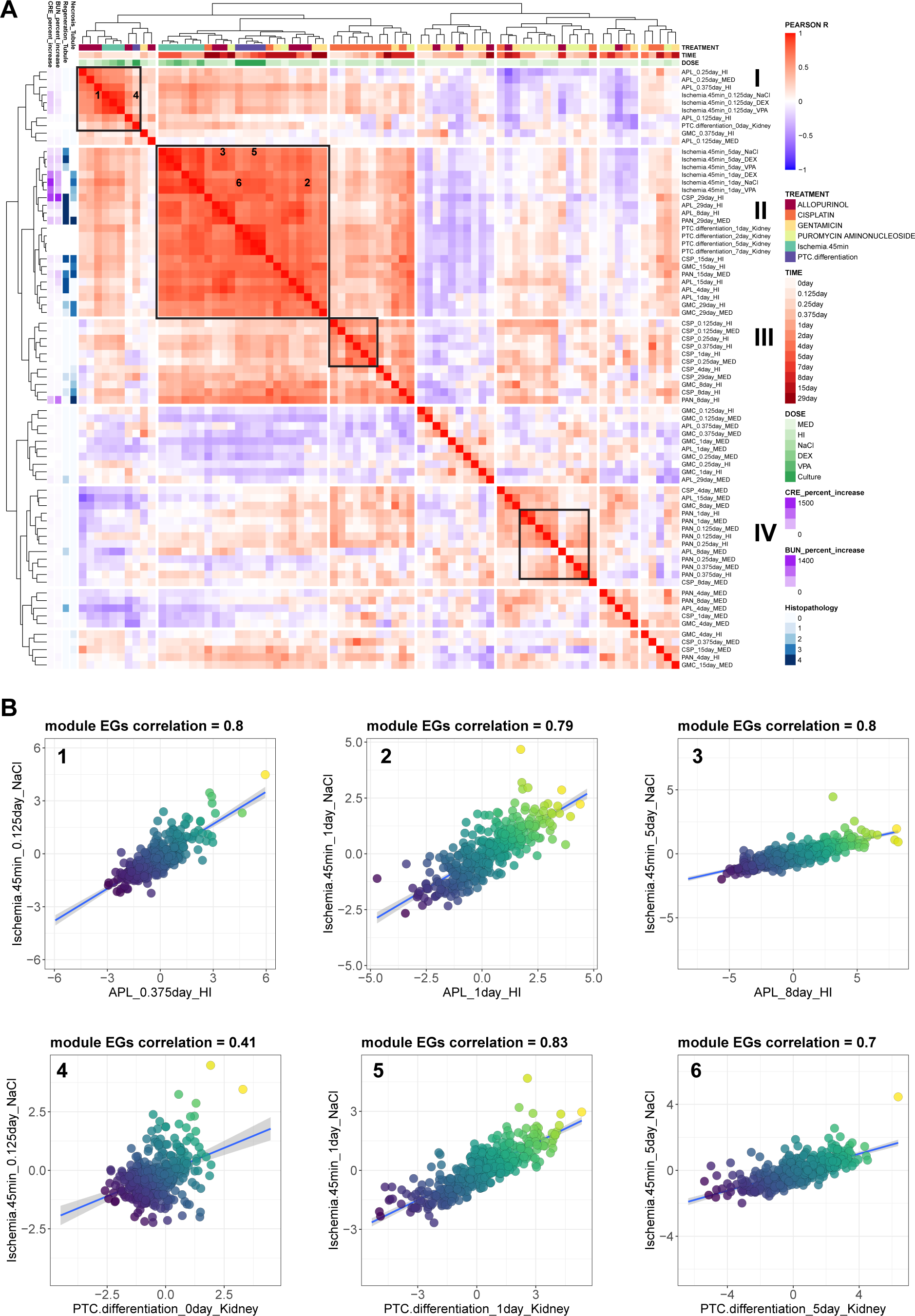
Correlation of ischemic injury and PTC dedifferentiation with nephrotoxic treatment conditions. (A) Cluster correlation plot for transcriptomic responses to nephrotoxic compounds, ischemic injury and primary PTC cultures, which shows strong Pearson R correlation between module EGS at time points when there is concurrent pathology (cluster II in black box), while early allopurinol (cluster I), cisplatin (cluster III) and puromycin aminonucleoside (cluster IV) responses are more distinct. Left heatmap displays the percent increase of serum injury biomarkers (BUN and CRE) compared to control (purple) and histopathology grades of tubule necrosis and regeneration (blue). Heatmap is clustered by Euclidean distance, using the complete method from *pheatmap* package. (B) Examples of correlation plots of responses during renal injury, showing high correlation between nephrotoxic treatments (APL = allopurinol), ischemic injury and PTC dedifferentiation.

To see if observations with nephrotoxicants can be extended to other form of renal injury, we analysed transcriptomic data from a previous rat renal ischemia reperfusion injury (IRI) study, a drug-independent model of acute kidney injury (AKI) ^23^. Gene expression data were uploaded into the TXG-MAPr environment and module EGS calculated (**Tables S11, S12**). The IRI study showed high similarity with early allopurinol treatment, mainly triggered by an early inflammatory response by both treatments (**Figures 4A cluster I, 4B panel 1, and S11A**). In contrast, the ischemic injury response at day 1 and 5 correlated well with late nephrotoxic TG treatments with co-occurring pathologies, suggesting that this is a general tissue response to injury, since there is no drug involved in vehicle (NaCl) treated ischemic kidneys (**Figures 4A cluster II, 4B panel 2, 3 and S11A**). Valproic acid (VPA) pre-exposure was able to accelerate the regeneration response at 1 day post ischemic injury compared to NaCl, an effect associated with a stronger activation of cell cycle module rKID:5m (**Figure S11B**).

Regenerating or injured proximal tubule epithelial cells (PTC) *in vivo* and in primary culture undergo dedifferentiation and regenerative responses, including a partial epithelial-to-mesenchymal transition, based on investigation of specific signalling pathways and expression of the mesenchymal intermediate filament, vimentin^52,53^. To determine if the global transcriptional response to dedifferentiation of cultured PTC bore any relationship to the dedifferentiation response associated with injury and regeneration *in vivo*, we compared rat PTC at various time in culture (*i.e.,* 1-7 days after plating on Transwell filters) with TG treatments (**Tables S11 and S12**). Notably, there was high similarity between the modular gene expression responses of cultured PTC and late nephrotoxic compound treatments (**Figures 4A cluster II, S11C and S12**), as well as to ischemic injury response at day 1 and 5, a time at which regeneration of proximal tubule epithelium is expected (**Figure 4A, 4B panel 4,5,6**). Thus, the dedifferentiation response of primary PTC appears similar, at the transcriptional level, to that of the regenerative PTC phenotype associated with the generalized tissue injury/regeneration response *in vivo*. Notably, activation of modules involved in cell cycle (rKID:5m), cytoskeleton / cell adhesion (rKID:2m, 42m), transcription (rKID:29, 64m), RNA and protein processing peaked in the first two days of culture compared to freshly isolated PTC (**Figure S12**). In addition, there was clear downregulation of metabolism, mitochondrial function and transport in cultured primary PTC (**Figure S12**), which was also seen after kidney injury *in vivo*.

In conclusion, high similarities between cellular and tissue responses during nephrotoxic treatments, ischemic injury and cultured primary PTC suggest that these are generalizable compound agnostic tissue responses to injury, dedifferentiation and repair. Responses to nephrotoxic compounds at early timepoints are more distinct and likely indicate the unique compound specific mode of action that could lead to cellular stress and later renal injury.

### Preservation analysis of rat kidney modules reveal common biological processes, stress responses and kidney specific networks

One of the most important safety decisions when compounds transition from nonclinical to clinical testing is whether hazards identified in nonclinical studies will translate to human safety concerns. Ideally, quantitative metrics could be used to determine the likelihood that biological events (*e.g.*, molecular initiating events (MIE) or key events (KE) in an adverse outcome pathway (AOP) framework) will translate across species. A unique aspect of network models, like WGCNA, is that preservation statistics can be applied to determine if (node-edge relationships in) biological networks are preserved across datasets or systems ^27^. We first tested robustness of network preservation statistics by comparing rat kidney modules based on all data versus the individual TG or DM rat kidney transcriptomic data separately using the Z-summary module preservation statistic ^27^. Notably, all modules are moderately preserved (274 modules = 79% with Z-summary > 2) or highly preserved (73 modules = 21% with Z-summary > 10) in the DM data (**Figure 5A, Table S10**). Self-preservation (testing for preservation in the dataset from which modules were derived) in DM is only slightly lower compared to the self-preservation of the larger TG dataset. In addition, there is good correlation between Z-summary scores for preserved modules in DM or TG self-preservation, including a subset of 67 modules that are highly preserved using both comparisons (**Figure 5B**, green line = Z-summary > 10). This preservation analysis suggests that both TG and DM rat kidney datasets capture similar gene co-expression patterns and biological responses.

**Figure 5.**
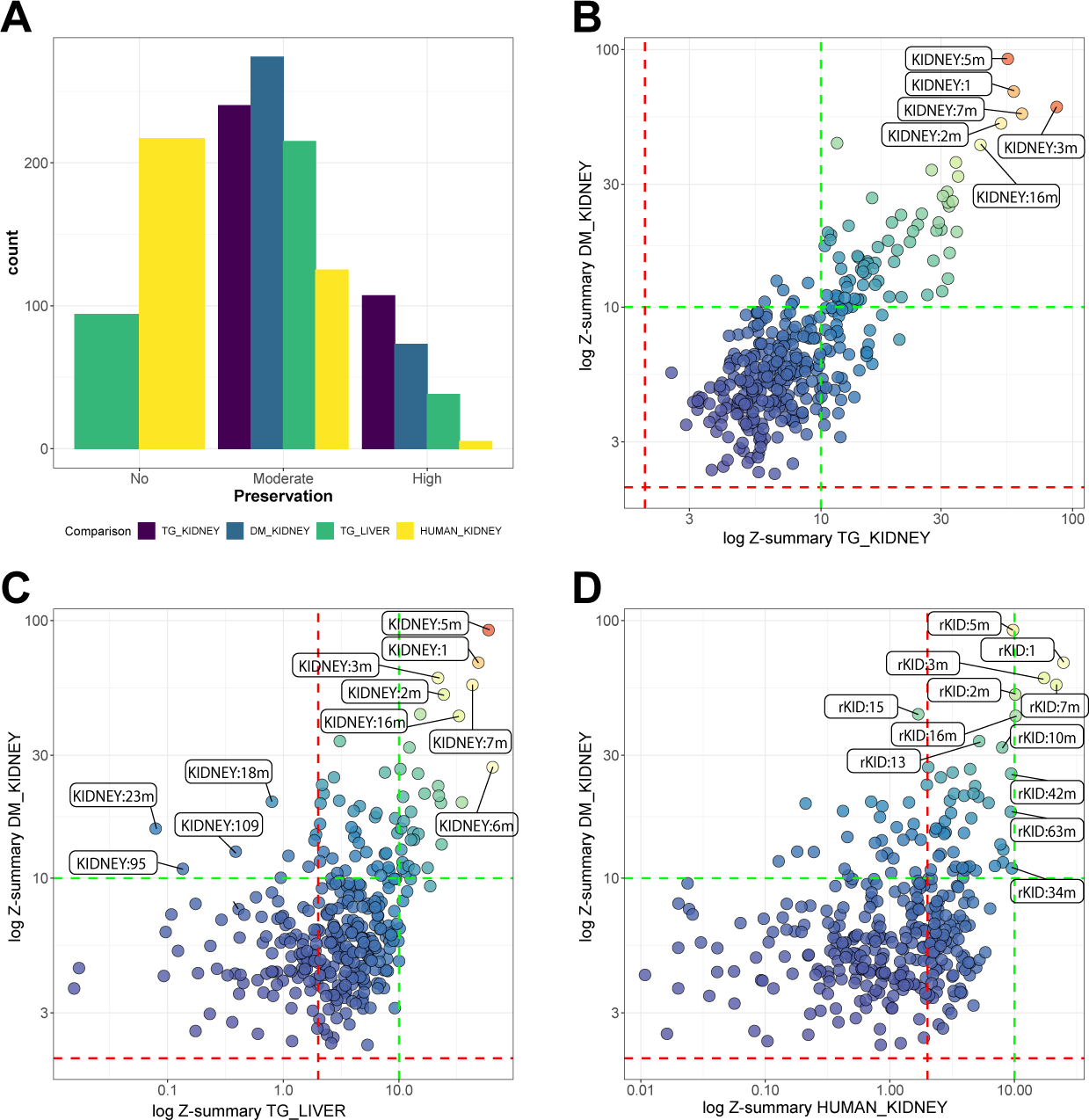
Module preservation. (A) Number of preserved rat kidney modules across datasets (TG-GATEs (TG) rat kidney, DrugMatrix (DM) rat kidney, TG rat liver and human kidney). TG and DM kidney datasets were used as a control to check if all modules are preserved. Z-summary > 2 indicate moderate module preservation and Z-summary > 10 indicate high module preservation. (B) Strong correlation between TG and DM preserved modules indicated by Z-summary scores for both comparisons, displayed on the axis as a log scale. Specific modules are highlighted that are strongly preserved in both datasets. (C) Comparison of rat kidney and liver Z-summary scores displayed on the axis as a log scale. Several modules are strongly preserved in both kidney and liver datasets, indicating that these processed have similar co-expression networks between the organs. Various kidney specific processes are only preserved in the kidney, but not in the liver (left of the red dashed line), including modules rKID:23m (kidney development), steroid metabolism (rKID:18m and 95) and Golgi function (rKID:109). (D) Comparison of rat kidney and human kidney Z-summary scores displayed on the axis as a log scale. Correlation plot of rat kidney modules preserved in DM kidney versus human kidney data, indicating that several modules are also preserved in human (rKID:2m, 3m, 5m, 7m, 10m, 16m and 42m amongst others), but others lack preservation in human (rKID:15 amongst others). Dashed red line indicates low/moderate module preservation threshold (Zsum > 2) and dashed green line indicates strong module preservation threshold (Zsum > 10). See also **Table S10** for more details.

To determine if highly preserved modules reflect common biological processes, we investigated their biological annotations. Highly preserved module annotate for processes important for maintaining cellular homeostasis as well as adaptive responses to chemical injury, including: cell cycle (rKID:5m), metabolism (rKID:3m, 9m, 14m, 21, 22, 25m, 116), mitochondrial function (rKID:26m, 28m, 37m, 43m, 96m), immune responses (rKID:7m, 15, 16m, 56, 63m, 113, 138, 212m), injury biomarker (rKID:2m), cytoskeleton (rKID:42m, 112, 142m), extracellular matrix (ECM) remodelling (rKID:10m, 31m), transport (rKID:13, 19, 78, 171), development (rKID:12m, 23m, 45m, 57m, 65, 70), ribosomal biogenesis / RNA processing (rKID:6m, 8m, 34m, 37m, 40m), proteasome (rKID:35m), circadian regulation (rKID:33) and responses to oxidative (rKID:53 = Nrf2) or endoplasmic reticulum (ER) stress (rKID:81 = Atf6; rKID:74 = Atf4). Interestingly, there was a large group of 255 kidney modules also preserved in rat liver TG-GATEs dataset (**Figure 5A and 5C, Table S10**), including the aforementioned modules involved in regulation of cellular stress or homeostasis. In contrast, some kidney specific modules were not preserved in liver, including modules involved in (renal) development (rKID:23m, 45m, 57m, 70, 186, 196, 290), steroid hormone metabolism (rKID:18m, 95) and transmembrane transport (rKID:171, 211, 329) amongst others (**Figure 5C**). Thus, preservation analysis illustrates that modules are highly preserved and robust reflections of cellular responses to drug-or chemical-induced renal injury, suggesting that the rat kidney TXG-MAPr analysis framework can be applied to other kidney toxicogenomic data to investigate DIKI mechanisms.

### Preserved injury and repair related networks in human patients

Having established that preservation of rat kidney modules is robust when comparing different large transcriptomics datasets of the same organ and species, we tested preservation between rat and human using available human kidney transcriptomic data. We evaluated preservation of rat kidney modules in a large human dataset (GEO: GSE36059) consisting of over 400 renal transplant samples from human patients at various times after transplantation with or without evidence of graft rejection as well as ‘control’ nephrectomy biopsies ^18^. Although the human dataset is limited to renal transplant samples, among the 347 rat kidney modules, 5 modules had strong preservation and 125 had moderate preservation in human kidneys (**Figure 5A**, **Table S10**). Modules involved in immune response (rKID:7m, rKID:16m), cell cycle (rKID:5m) and metabolism (rKID:3m) were strongly preserved (**Figure 5D**), while several other modules representing other immune responses, metabolism, mitochondrial function, RNA processing, cell cycle and ECM remodelling/fibrosis were moderately preserved (**Table S10**). These observations are in line with the fact that a subset of the human samples were patients experiencing immune response-mediated graft rejection ^18,24^. Strikingly, the cytoskeleton or injury biomarker module rKID:2m was also strongly preserved in human, showing the relevance of the biological context of the renal biomarkers that cluster in similar co-expressed modules (**Figure 5D**, **Table S10**). Other potential biomarker modules with strong pathology association were also preserved in human (**Tables 1, 2 and S10**). By contrast, some of the modules were not (or hardly) preserved (Zsum ≤ 2) in the available human dataset, including modules reflecting cellular stress responses, like ATF4 (rKID:74), oxidative stress (rKID:53, 105, 106), as well as modules annotated for ribosomal biogenesis (rKID:6m, 8m), renal development (rKID:23m, 45m), circadian regulation (rKID:33), immune response (rKID:15, 56), steroid hormone metabolism (rKID:18m, 95), and several other metabolic processes (**Table S10**).

Next, the human dataset was uploaded into the rat kidney TXG-MAPr tool (after gene ortholog conversion) to see how the human responses were captured using only the preserved modules (**Figures 6 and S13, Table S13**). As expected, immune responses (rKID:7m, 16m, 63m, 113, 138, 212m 305), extracellular matrix (rKID:10m, 31m 104) and cell migration (rKID:127) modules were most strongly induced in human transplant samples and clustered together in the heatmap (**Figure S13**, cluster 3). More inter-individual variation was seen in modules involved in mitochondrial function, metabolism and transport, which grouped together in cluster 2 (**Figure S13**). Immune response and injury biomarker modules showed on average stronger activation in patients with immune-mediated renal rejection despite some inter-individual variation, while metabolism was stronger downregulated in these patients (**Figure 6A, S14**). Similar patterns could be seen for the log2FC of hub genes and most significant genes of each of these inflammation modules (**Figure 6A, S14**). In addition, inflammation modules rKID:7m and rKID:63m showed a very strong correlation (Pearson R = 0.97) across all patient samples (**Figure 6B, left**), which was similar for most of the inflammation and cell migration modules (**Figure 6C, cluster 3**). Also, the biomarker module rKID:2m showed strong correlation (Pearson R = 0.72) with inflammation module rKID:7m (**Figure 6B, centre**). In contrast, metabolism module rKID:3m showed inverse correlation (Pearson R = −0.62) with inflammation module rKID:7m (**Figure 6B, right**) and rKID:2m, similar to responses in rat kidney. This bi-directional response was also seen for other metabolism, mitochondrial and transport modules compared to inflammation (**Figure 6C, cluster 1 vs 3**). This indicates that metabolism, mitochondrial function and transport are down regulated in patients’ kidneys undergoing renal inflammation, which was also noted in rats. A large group of modules with various annotations for RNA and protein processing showed very high correlation (**Figure 6, cluster 4**), which was mainly due to downregulation of these modules in a group of patients with highest AvAbsEGS. With the presented gene co-expression approach in pre-clinical models, we could identify several modules that were preserved in human, indicating the relevance for human translation. Moreover, we showed differential gene network dynamics in human patients with rejected renal grafts that could be linked to immune responses, injury and biomarker expression, indicating that human and rat have similar generalized tissue responses to kidney injury and regeneration.

**Figure 6.**
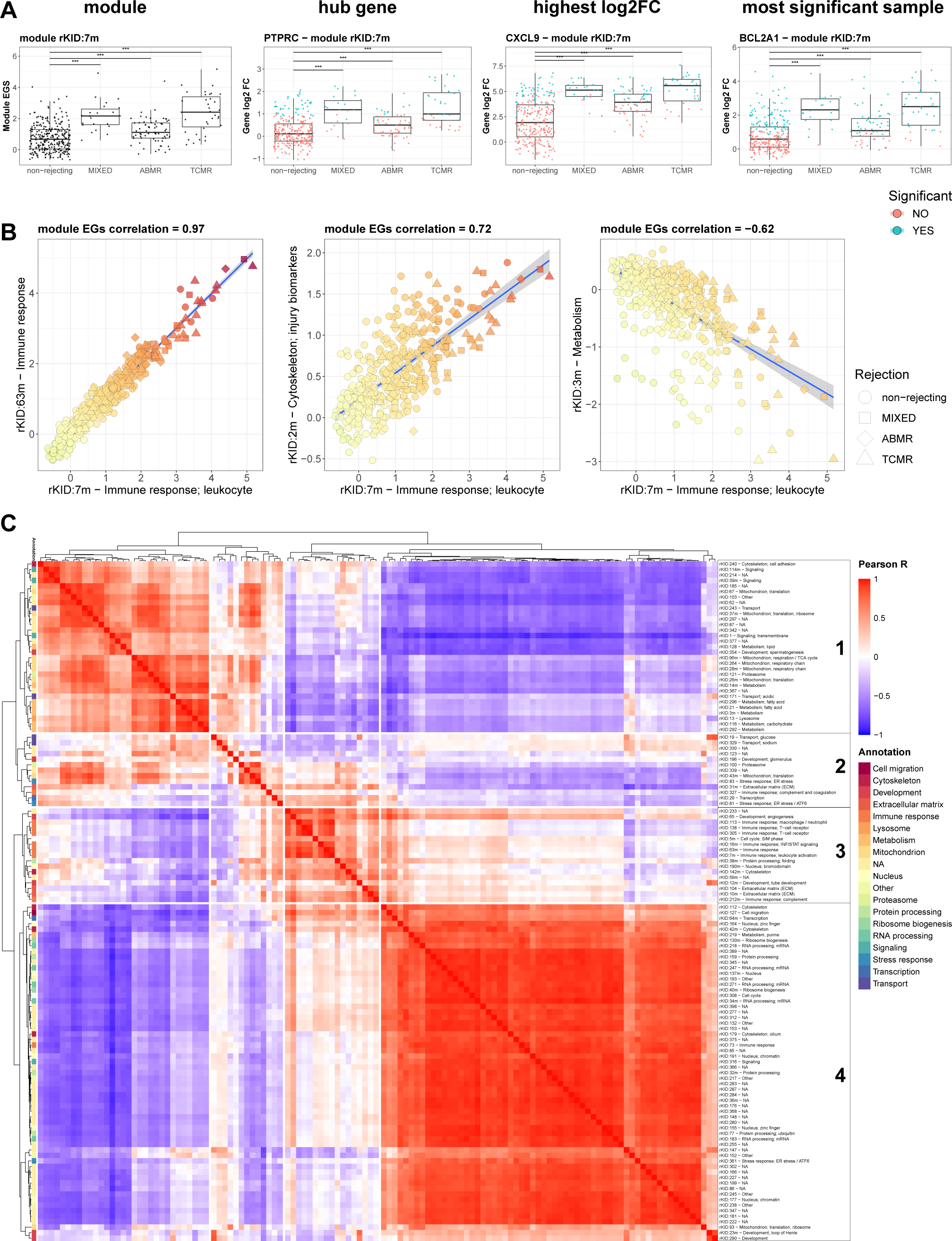
Human preservation and network dynamics. (A) Box plot for the different human transplant groups showing the module EGS (left) and the log2 fold change of the hub gene (second column) and most significant genes of inflammation module rKID:7m. The groups are indicated by rejection type: non-rejecting, ABMR = antibody mediated rejection, TCMR = T-cell mediated rejection, MIXED = both antibody and T-cell mediated rejection. Significant differences between groups were calculated using a Wilcoxon test. Significant gene log2 fold changes of individual patients were indicated in blue (p < 0.05). (B) Module EGS correlation plot of inflammation modules rKID:7m vs rKID:63m (left), inflammation module rKID:7m vs injury biomarker module rKID:2m (centre) and inflammation rKID:7m vs metabolism rKID:3m modules. Strong correlation is shown between the two inflammation and injury biomarker modules, while there is inverse correlation between the inflammation module rKID:7m and the metabolism module rKID:3m. (C) Cluster correlation heatmap (based on Pearson correlation of module EGS) of the preserved modules in human with a EGS > 2. There is strong clustering of the inflammation, ECM and cell migration modules (cluster 3) and cellular stress or injury related modules (cluster 4). The mitochondrial function, metabolism and transport modules cluster together as well (cluster 1) but show anti correlation with the inflammation and stress / injury modules. Heatmap is clustered by Euclidean distance, using the complete method from *pheatmap* package.

## Discussion

In pre-clinical research a wealth of toxicogenomic data has been generated in the past decades, with the aim to investigate biological responses to drugs and to understand the mechanisms of toxicity ^7^. Unfortunately, while standardized reporting approaches for bioinformatics have been proposed ^54–56^, a comprehensive qualitative and quantitative mechanistic interpretation framework for toxicogenomics underpinning a mechanism-based safety assessment approach is needed ^7,8^. A major hurdle has been the data complexity and the inability to generalize conclusions from transcriptomics responses across different cell types, tissues and species. Methods that reduce complex transcriptomic data to networks of co-expressed genes that retain biological meaning are an attractive approach to reducing the dimensionality of transcriptomic data and facilitating interpretation ^15,57^, as demonstrated previously for rat liver toxicity ^10–13^. Besides liver injury, kidney toxicity is also a common adverse response in both drug and chemical toxicity ^1^. Therefore, methods that facilitate the extraction of mechanistic information from kidney transcriptional responses would be useful in hazard identification and risk assessment.

Herein, we used WGCNA to construct a set of 347 co-regulated gene networks (modules) from the two largest, publicly available *in vivo* rat kidney toxicogenomics datasets (TG-GATEs and DrugMatrix) and provide biological annotations for the modules allowing direct translation of gene expression data into meaningful biological conclusions. The modules are embedded in an interactive R-shiny application, called the rat kidney TXG-MAPr, that allows dose- and time-response analyses, treatment correlation and functional annotation (GO-terms, pathways and transcription factor enrichment). We demonstrated that this gene co-expression approach and analysis framework has the advantage that modules and their quantitative EGS can assist in: (1) data complexity reduction, (2) biological annotation for mechanistic interpretation, (3) pathology association, (4) biomarker discovery, and (5) translation to human relevant responses.

First, we have demonstrated that the co-expression modules could identify meaningful biological responses relevant to mechanistic investigation using cisplatin exposure as the example. Early activation of DNA-damage and p53 signalling (rKID:160) was followed by later occurrence of cellular injury and biomarker expression (rKID:2M), regeneration (rKID:5m) and inflammation (rKID:56, 63m, 212m and 327). These later responses, including activation of biomarker-containing, regeneration and inflammatory modules, were mirrored with other nephrotoxicants that had concurrent renal histopathology and clinical chemistry changes. This suggests that similarity in gene expression patterns reflects common injury-induced repair mechanisms (*e.g.*, regeneration, dedifferentiation and inflammation responses). The common injury and regeneration processes across different nephrotoxicants, as well as during ischemic injury, are compound agnostic and general tissue responses to injury and repair, where a complex interplay between renal inflammation and cellular dedifferentiation or partial EMT are contributing to renal repair ^58^. However, at early stages after nephrotoxic compounds exposure there was a distinct activation of biological response networks relevant to the expected different mechanisms of injury. For example, the induction of rKID:160 and rKID:298 (the modules annotating for p53 responses) is much stronger and unique for cisplatin, carboplatin and doxorubicin, due to their genotoxic mechanism of action, compared to other nephrotoxicants, like puromycin aminonucleoside. In contrast, the primary key event of puromycin is an early inflammatory response, which the anticipated mechanisms of action that can induce glomerular nephropathy ^50^. In addition, estrogen receptor agonists are the main inducers of steroid hormone metabolism modules, which is expected based on their primary mechanism of action. Thus, the rat kidney TXG-MAPr tool also allows evaluation of molecular mechanisms and early mode-of-action of drugs and chemicals with respect to onset of nephrotoxicity. The module annotations that we provide in this study enhance a qualitative and quantitative mechanistic interpretation of renal toxicogenomics data and capture most of the well-known biological processes.

Various studies have applied gene co-expression network analysis on rat and human kidney injury samples ^59–64^. A previous study applied gene co-expression analysis on the rat kidney DM data and also identified modules that correlated with acute kidney injury, which contained *Havcr1* and *Clu* genes ^59^, which are in module rKID:2m with the strongest pathology association in our study as well. Most other studies identified a small number of modules due to the limited number of samples used to build the gene co-expression networks, but the captured modules include stress and immune responses, which were also the most relevant modules for pathology in our study. However, these previous studies did not integrate their kidney gene co-expression networks into a publicly accessible online tool, like the rat kidney TXG-MAPr, underpinning its value. Though there are co-expression tools built from data of various species and tissues that are publicly available ^65–69^. These tools mainly show gene interactions, pathway analysis, or conservation across species. In contrast, the rat kidney TXG-MAPr tool allows the users to interactively visualize quantitative changes in gene networks by chemical or biological perturbations. In addition, users can upload log2FC gene expression data for comparison to the reference toxicogenomics dataset in the TXG-MAPr tool, which is highly valuable to identify common mechanisms of action for chemical hazard identification.

Inflammation accompanied by metabolic re-wiring are hallmarks of acute tubular injury and the balance between injury and repair ^51^. Thus, the juxtaposition of inflammation module induction and repression of metabolism and mitochondrial modules is worth noting. Repression of module rKID:3m (highly annotated for small molecular metabolism and mitochondrion (**Table S3**)) ranked among the top ten modules (based on effect size and p-adjust) for multiple pathologies (**Tables S8 and S9**). Conversely, modules rKID:7m, 63m and 327 are annotated for immune response and were among the top five ranked inducible modules associated with the presence of cellular infiltrates, *i.e.,* inflammation (**Tables 2 and S8**). Thus, strong activation in regeneration and inflammation responses impacted on cellular energy production (metabolism and mitochondrial function) in the kidney, which was also concluded previously for liver diseases ^70^. The bi-directional response of these processes was also demonstrated in human patients with renal transplants (**Figure 6C**), suggesting that these injury and repair responses are also conserved between species.

In this study we have identified modules with strong statistical association with histopathology findings and biomarkers of tissue injuries. Module rKID:2m was most strongly associated with concurrent and predictive renal pathology and contained many well-known kidney injury biomarkers, including *Havcr1* (Kim-1), *Lcn2* (Lipocalin 2) *Spp1* (Osteopontin) and *Clu* (Clusterin). In addition, several immune response modules ranked highly for association with concurrent and predictive kidney injury (**Tables 2 and S9**), including modules rKID:327, rKID:56 and rKID:212m, containing hub genes that are candidate novel biomarkers such as fibrinogen (*Fga, Fgb, Fgg*), *Icam1* and ceruloplasmin (*CP*), respectively ^71–74^. This suggests that novel injury biomarkers could be discovered using gene co-expression networks that are strongly associated with pathology, as was recently reported for liver and kidney ^11,75^. This approach was validated in a recently published study where early activation of injury modules after cisplatin exposure will lead to renal pathology in rats by 28 days, including necrosis, inflammation and fibrosis ^76^. It was shown that urinary clusterin correlated well with *Clu* gene expression, which is an important biomarker in rKID:2m that showed the strongest pathology association. The subset of both concurrent and predictive kidney injury-associated modules is important for chemical or drug development since aberrant transcriptional activity could indicate higher risk of developing adversity. Moreover, association of (early) WGCNA module changes that are predictive of late-stage kidney pathology is expected to impact drug candidate optimization and enable prioritization of safer compounds.

Preservation analysis is a unique feature of network-based models allowing not only the robustness and reproducibility of modules to be tested using datasets from the same experimental species, but also the likelihood of translation across species, which is an important consideration in drug or chemical hazard assessment. All rat kidney modules had at least moderate self-preservation in the TG and DM rat kidney transcriptomic data, indicating that the modules were robust and reproducible representations of biological response networks in rat kidneys. However, only 130 (37%) of the 347 rat modules were preserved (translational modules) when compared to a human renal transplantation transcriptomic dataset. Among the translational modules were those that reflect immune response, cell cycle, metabolism, mitochondrial function, RNA processing, and ECM remodelling / fibrosis modules. These biological programs are deterministic of end-stage renal disease, involving inflammatory and regenerative programs that are part of renal transplant rejection ^51^. By contrast, various modules that reflect early cellular stress response programs, including ER stress activation of Atf4, Nrf2-mediated oxidative stress, as well as modules annotated for ribosomal biogenesis, renal development, circadian regulation, steroid hormone metabolism were not preserved in human. These differences are likely due to the nature of the human dataset, which only contained human transplant samples, where early injury or disease responses, including adaptive cellular stress responses, are not captured in these late-stage human transplant disease settings ^18^. Nonetheless, the fact that important toxicity-related rat kidney modules are preserved in human kidney samples suggests that the preservation approach is an attractive and valid strategy to determine the likelihood that mechanisms of renal injury observed in rat kidney will translate to human. Future studies should focus on collection of more extensive and diverse human renal pathology transcriptomics datasets representing a broader spectrum of early and late-stage disease phenotypes, which may improve the preservation statistics and confidence which rat modules will translate to human, which is significant knowledge for chemical safety assessment.

In conclusion, the rat kidney TXG-MAPr tool provides a user-friendly interface that enables visualization of gene expression data in the context of co-expressed modules that can be mined for the association of important biological processes in acute and chronic renal injury. The tool can be utilized for identifying possible safety liabilities and/or mechanisms that can lead to adversity, useful for chemical safety assessment. Association of early WGCNA module changes that are predictive of later kidney pathology is expected to impact lead optimization and enable prioritization of better compounds which are less likely to induce pathology. We foresee that quantitative gene co-expression modules, which are strongly associated to kidney pathology, and which translate from rat to human, could potentially be implemented in translational quantitative systems toxicology models for mechanistic evaluation and prediction of kidney injury of new chemical or drug candidates ^77^.

## Acknowledgements

This work has received funding from the EU-EFPIA Innovative Medicines Initiative 2 (IMI2) Joint Undertaking TransQST project (grant number 116030) and eTRANSAFE project (grant number 777365) (This Joint Undertaking receives support from the European Union’s Horizon 2020 research and innovation program and EFPIA), the EC Horizon2020 EU-ToxRisk project (grant number 681002), the EC Horizon2020 RISK-HUNT3R project (grant number 964537), the Horizon Europe PARC programme (The European Partnership for the Assessment of Risks from Chemicals, grant number 101057014) and the Virtual Human Platform for safety assessment (VHP4SAFETY) project (grant number 1292.19.272, which is part of the NWA research program ‘Research along Routes by Consortia (ORC)’, which is funded by the Netherlands Organization for Scientific Research (NWO).

Authors declare that this work reflects only the author’s view and that the European Commission and IMI-JU are not responsible for any use that may be made of the information it contains.

## Author contributions

S.J.K., G.C., J.J.S. and P.T. performed data analysis. S.J.K., G.C. and H.W.v.K. built the R-shiny TXG-MAPr tool. S.J.K., L.S.W., Git C., K.P., C.B., K.M.G., C.R.T., J.L.S., B.v.d.W. designed experiments. S.J.K., Git C., K.P. and K.M.G. performed experiments. G.C., J.J.S., P.T., C.P.F., J.S.R., C.B., S.A.E. and K.M.H.T. provided expertise and feedback. S.J.K., J.L.S. and B.v.d.W. conceptualized, wrote and edited the manuscript. J.L.S. and B.v.d.W. acquired research funding and supervised the study. All authors approved the manuscript.

## Abbreviations

AKI: Acute kidney injury
APL: Allopurinol
ATF: Activating transcription factor
AvAbsEGS: Average absolute eigengene score
BUN: Blood urea nitrogen
corEG: Correlation eigengene score
CRE: Creatinine
CSP: Cisplatin
DIKI: Drug-induced kidney injury
DM: DrugMatrix
EGS: Eigengene score
ER: Endoplasmic reticulum
FC: Fold change
GO: Gene ontology
IRI: Ischemia reperfusion injury
KIM-1: Kidney injury molecule-1
log2FC: log2 fold change
NRF2: Nuclear Factor Erythroid 2-Related Factor 2
ORA: Over Representation Analysis
PAN: Puromycin aminonucleoside
PTC: Proximal tubular cells
RMA: Robust Multi-array Average
TG: TG-GATEs
TF: Transcription factor
TXG: Toxicogenomics
WGCNA: Weighted gene co-expression network analysis

## References

1. Perazella, M.A. (2019). Drug-induced acute kidney injury: Diverse mechanisms of tubular injury. Curr Opin Crit Care 25, 550–557. 10.1097/MCC.0000000000000653.

2. Rayego-Mateos, S., Marquez-Expósito, L., Rodrigues-Diez, R., Sanz, A.B., Guiteras, R., Doladé, N., Rubio-Soto, I., Manonelles, A., Codina, S., Ortiz, A., et al. (2022). Molecular Mechanisms of Kidney Injury and Repair. Int J Mol Sci 23, 1542. 10.3390/ijms23031542.

3. Perazella, M.A. (2018). Pharmacology behind common drug nephrotoxicities. Clinical Journal of the American Society of Nephrology 13, 1897–1908. 10.2215/CJN.00150118.

4. Irazabal, M. V., and Torres, V.E. (2020). Reactive Oxygen Species and Redox Signaling in Chronic Kidney Disease. Cells 9, 1342. 10.3390/cells9061342.

5. Yan, M., Shu, S., Guo, C., Tang, C., and Dong, Z. (2018). Endoplasmic reticulum stress in ischemic and nephrotoxic acute kidney injury. Ann Med 50, 381–390. 10.1080/07853890.2018.1489142.

6. McDuffie, J.E. (2018). Brief Overview: Assessment of Compound-induced Acute Kidney Injury Using Animal Models, Biomarkers, and In Vitro Platforms. Toxicol Pathol 46, 978–990. 10.1177/0192623318807679.

7. Vahle, J.L., Anderson, U., Blomme, E.A.G., Hoflack, J.C., and Stiehl, D.P. (2018). Use of toxicogenomics in drug safety evaluation: Current status and an industry perspective. Regulatory Toxicology and Pharmacology 96, 18–29. 10.1016/j.yrtph.2018.04.011.

8. Pettit, S., des Etages, S.A., Mylecraine, L., Snyder, R., Fostel, J., Dunn, R.T., Haymes, K., Duval, M., Stevens, J., Afshari, C., et al. (2010). Current and Future Applications of Toxicogenomics: Results Summary of a Survey from the HESI Genomics State of Science Subcommittee. Environ Health Perspect 118, 992–997. 10.1289/ehp.0901501.

9. Ideker, T., Dutkowski, J., and Hood, L. (2011). Boosting Signal-to-Noise in Complex Biology: Prior Knowledge Is Power. Cell 144, 860–863. 10.1016/j.cell.2011.03.007.

10. Callegaro, G., Kunnen, S.J., Trairatphisan, P., Grosdidier, S., Niemeijer, M., den Hollander, W., Guney, E., Piñero Gonzalez, J., Furlong, L., Webster, Y.W., et al. (2021). The human hepatocyte TXG-MAPr: gene co-expression network modules to support mechanism-based risk assessment. Arch Toxicol 95, 3745– 3775. 10.1007/s00204-021-03141-w.

11. Callegaro, G., Schimming, J.P., Piñero González, J., Kunnen, S.J., Wijaya, L., Trairatphisan, P., van den Berk, L., Beetsma, K., Furlong, L.I., Sutherland, J.J., et al. (2023). Identifying multiscale translational safety biomarkers using a network-based systems approach. iScience 26, 106094. 10.1016/j.isci.2023.106094.

12. Sutherland, J.J., Jolly, R.A., Goldstein, K.M., and Stevens, J.L. (2016). Assessing Concordance of Drug-Induced Transcriptional Response in Rodent Liver and Cultured Hepatocytes. PLoS Comput Biol 12. 10.1371/journal.pcbi.1004847.

13. Sutherland, J.J., Webster, Y.W., Willy, J.A., Searfoss, G.H., Goldstein, K.M., Irizarry, A.R., Hall, D.G., and Stevens, J.L. (2018). Toxicogenomic module associations with pathogenesis: a network-based approach to understanding drug toxicity. Pharmacogenomics J 18, 377–390. 10.1038/tpj.2017.17.

14. Zhang, B., and Horvath, S. (2005). A general framework for weighted gene co-expression network analysis. Stat Appl Genet Mol Biol 4. 10.2202/1544-6115.1128.

15. Langfelder, P., and Horvath, S. (2008). WGCNA: An R package for weighted correlation network analysis. BMC Bioinformatics 9, 559. 10.1186/1471-2105-9-559.

16. Igarashi, Y., Nakatsu, N., Yamashita, T., Ono, A., Ohno, Y., Urushidani, T., and Yamada, H. (2015). Open TG-GATEs: A large-scale toxicogenomics database. Nucleic Acids Res 43, D921--D927. 10.1093/nar/gku955.

17. Svoboda, D.L., Saddler, T., and Auerbach, S.S. (2019). An Overview of National Toxicology Program’s Toxicogenomic Applications: DrugMatrix and ToxFX. In, Hong H., ed. (Springer), pp. 141–157. 10.1007/978-3-030-16443-0_8.

18. Reeve, J., Sellarés, J., Mengel, M., Sis, B., Skene, A., Hidalgo, L., De Freitas, D.G., Famulski, K.S., and Halloran, P.F. (2013). Molecular diagnosis of T cell-mediated rejection in human kidney transplant biopsies. American Journal of Transplantation 13, 645–655. 10.1111/ajt.12079.

19. R Core Team (2024). R: A Language and Environment for Statistical Computing. Preprint.

20. Gautier, L., Cope, L., Bolstad, B.M., and Irizarry, R.A. (2004). affy--analysis of Affymetrix GeneChip data at the probe level. Bioinformatics 20, 307–315. 10.1093/bioinformatics/btg405.

21. Dai, M., Wang, P., Boyd, A.D., Kostov, G., Athey, B., Jones, E.G., Bunney, W.E., Myers, R.M., Speed, T.P., Akil, H., et al. (2005). Evolving gene/transcript definitions significantly alter the interpretation of GeneChip data. Nucleic Acids Res 33, e175–e175. 10.1093/nar/gni179.

22. Ritchie, M.E., Phipson, B., Wu, D., Hu, Y., Law, C.W., Shi, W., and Smyth, G.K. (2015). limma powers differential expression analyses for RNA-sequencing and microarray studies. Nucleic Acids Res 43, e47– e47. 10.1093/nar/gkv007.

23. Speir, R.W., Stallings, J.D., Andrews, J.M., Gelnett, M.S., Brand, T.C., and Salgar, S.K. (2015). Effects of Valproic Acid and Dexamethasone Administration on Early Bio-Markers and Gene Expression Profile in Acute Kidney Ischemia-Reperfusion Injury in the Rat. PLoS One 10, e0126622. 10.1371/journal.pone.0126622.

24. Halloran, P.F., Chang, J., Famulski, K., Hidalgo, L.G., Salazar, I.D.R., Lopez, M.M., Matas, A., Picton, M., De Freitas, D., Bromberg, J., et al. (2015). Disappearance of T cell-mediated rejection despite continued antibody-mediated rejection in late kidney transplant recipients. Journal of the American Society of Nephrology 26, 1711–1720. 10.1681/ASN.2014060588.

25. Brown, C.D.A., Sayer, R., Windass, A.S., Haslam, I.S., De Broe, M.E., D’Haese, P.C., and Verhulst, A. (2008). Characterisation of human tubular cell monolayers as a model of proximal tubular xenobiotic handling. Toxicol Appl Pharmacol 233, 428–438. 10.1016/j.taap.2008.09.018.

26. Bajaj, P., Chung, G., Pye, K., Yukawa, T., Imanishi, A., Takai, Y., Brown, C., and Wagoner, M.P. (2020). Freshly isolated primary human proximal tubule cells as an in vitro model for the detection of renal tubular toxicity. Toxicology 442, 152535. 10.1016/j.tox.2020.152535.

27. Langfelder, P., Luo, R., Oldham, M.C., and Horvath, S. (2011). Is my network module preserved and reproducible? PLoS Comput Biol 7, 1001057. 10.1371/journal.pcbi.1001057.

28. Alexa, A., Rahnenführer, J., and Lengauer, T. (2006). Improved scoring of functional groups from gene expression data by decorrelating GO graph structure. Bioinformatics 22, 1600–1607. 10.1093/bioinformatics/btl140.

29. Kamburov, A., Stelzl, U., Lehrach, H., and Herwig, R. (2013). The ConsensusPathDB interaction database: 2013 Update. Nucleic Acids Res 41. 10.1093/nar/gks1055.

30. Garcia-Alonso, L., Holland, C.H., Ibrahim, M.M., Turei, D., and Saez-Rodriguez, J. (2019). Benchmark and integration of resources for the estimation of human transcription factor activities. Genome Res 29, 1363–1375. 10.1101/gr.240663.118.

31. Alvarez, M.J., Shen, Y., Giorgi, F.M., Lachmann, A., Ding, B.B., Hilda Ye, B., and Califano, A. (2016). Functional characterization of somatic mutations in cancer using network-based inference of protein activity. Nat Genet 48, 838–847. 10.1038/ng.3593.

32. Chang, W., Cheng, J., Allaire, J.J., Sievert, C., Schloerke, B., Xie, Y., Allen, J., McPherson, J., Dipert, A., and Borges, B. (2024). shiny: Web Application Framework for R. Preprint.

33. Wickham, H. (2016). ggplot2: Elegant Graphics for Data Analysis (Springer-Verlag New York).

34. Kolde, R. (2019). pheatmap: Pretty Heatmaps. Preprint.

35. Manohar, S., and Leung, N. (2018). Cisplatin nephrotoxicity: a review of the literature. J Nephrol 31, 15–25. 10.1007/s40620-017-0392-z.

36. McWilliam, S.J., Wright, R.D., Welsh, G.I., Tuffin, J., Budge, K.L., Swan, L., Wilm, T., Martinas, I.-R., Littlewood, J., and Oni, L. (2021). The complex interplay between kidney injury and inflammation. Clin Kidney J 14, 780–788. 10.1093/ckj/sfaa164.

37. Rajan, V., and Mitch, W.E. (2008). Ubiquitin, proteasomes and proteolytic mechanisms activated by kidney disease. Biochim Biophys Acta Mol Basis Dis 1782, 795–799. 10.1016/j.bbadis.2008.07.007.

38. Ichii, O., Otsuka-Kanazawa, S., Nakamura, T., Ueno, M., Kon, Y., Chen, W., Rosenberg, A.Z., and Kopp, J.B. (2014). Podocyte Injury Caused by Indoxyl Sulfate, a Uremic Toxin and Aryl-Hydrocarbon Receptor Ligand. PLoS One 9, e108448. 10.1371/journal.pone.0108448.

39. Grgic, I., Hofmeister, A.F., Genovese, G., Bernhardy, A.J., Sun, H., Maarouf, O.H., Bijol, V., Pollak, M.R., and Humphreys, B.D. (2014). Discovery of new glomerular disease–relevant genes by translational profiling of podocytes in vivo. Kidney Int 86, 1116–1129. 10.1038/ki.2014.204.

40. Han, W.K., Bailly, V., Abichandani, R., Thadhani, R., and Bonventre, J. V. (2002). Kidney Injury Molecule-1 (KIM-1): A novel biomarker for human renal proximal tubule injury. Kidney Int 62, 237–244. 10.1046/j.1523-1755.2002.00433.x.

41. Dieterle, F., Sistare, F., Goodsaid, F., Papaluca, M., Ozer, J.S., Webb, C.P., Baer, W., Senagore, A., Schipper, M.J., Vonderscher, J., et al. (2010). Renal biomarker qualification submission: a dialog between the FDA-EMEA and Predictive Safety Testing Consortium. Nat Biotechnol 28, 455–462. 10.1038/nbt.1625.

42. Wen, Y., and Parikh, C.R. (2021). Current concepts and advances in biomarkers of acute kidney injury. Crit Rev Clin Lab Sci 58, 354–368. 10.1080/10408363.2021.1879000.

43. Teo, S.H., and Endre, Z.H. (2017). Biomarkers in acute kidney injury (AKI). Best Pract Res Clin Anaesthesiol 31, 331–344. 10.1016/j.bpa.2017.10.003.

44. Zhang, W.R., and Parikh, C.R. (2019). Biomarkers of Acute and Chronic Kidney Disease. Annu Rev Physiol 81, 309–333. 10.1146/annurev-physiol-020518-114605.

45. Srisawat, N., and Kellum, J.A. (2020). The Role of Biomarkers in Acute Kidney Injury. Crit Care Clin 36, 125–140. 10.1016/j.ccc.2019.08.010.

46. Vaidya, V.S., Ferguson, M.A., and Bonventre, J. V. (2008). Biomarkers of Acute Kidney Injury. Annu Rev Pharmacol Toxicol 48, 463–493. 10.1146/annurev.pharmtox.48.113006.094615.

47. Sieber, M., Hoffmann, D., Adler, M., Vaidya, V.S., Clement, M., Bonventre, J. V., Zidek, N., Rached, E., Amberg, A., Callanan, J.J., et al. (2009). Comparative Analysis of Novel Noninvasive Renal Biomarkers and Metabonomic Changes in a Rat Model of Gentamicin Nephrotoxicity. Toxicological Sciences 109, 336–349. 10.1093/toxsci/kfp070.

48. Huang, H., Lin, Q., Dai, X., Chen, J., Bai, Z., Li, X., Fang, F., and Li, Y. (2022). Derivation and validation of urinary TIMP-1 for the prediction of acute kidney injury and mortality in critically ill children. J Transl Med 20, 102. 10.1186/s12967-022-03302-0.

49. Tummalapalli, L., Nadkarni, G.N., and Coca, S.G. (2016). Biomarkers for predicting outcomes in chronic kidney disease. Curr Opin Nephrol Hypertens 25, 480–486. 10.1097/MNH.0000000000000275.

50. Petrovic-Djergovic, D., Popovic, M., Chittiprol, S., Cortado, H., Ransom, R.F., and Partida-Sánchez, S. (2015). CXCL10 induces the recruitment of monocyte-derived macrophages into kidney, which aggravate puromycin aminonucleoside nephrosis. Clin Exp Immunol 180, 305–315. 10.1111/cei.12579.

51. Tammaro, A., Kers, J., Scantlebery, A.M.L., and Florquin, S. (2020). Metabolic Flexibility and Innate Immunity in Renal Ischemia Reperfusion Injury: The Fine Balance Between Adaptive Repair and Tissue Degeneration. Front Immunol 11. 10.3389/fimmu.2020.01346.

52. Torsello, B., De Marco, S., Bombelli, S., Cifola, I., Morabito, I., Invernizzi, L., Meregalli, C., Zucchini, N., Strada, G., Perego, R.A., et al. (2023). High glucose induces an activated state of partial epithelial-mesenchymal transition in human primary tubular cell cultures. PLoS One 18, e0279655. 10.1371/journal.pone.0279655.

53. De Chiara, L., and Crean, J. (2016). Emerging Transcriptional Mechanisms in the Regulation of Epithelial to Mesenchymal Transition and Cellular Plasticity in the Kidney. J Clin Med 5, 6. 10.3390/jcm5010006.

54. Bourdon-Lacombe, J.A., Moffat, I.D., Deveau, M., Husain, M., Auerbach, S., Krewski, D., Thomas, R.S., Bushel, P.R., Williams, A., and Yauk, C.L. (2015). Technical guide for applications of gene expression profiling in human health risk assessment of environmental chemicals. Regulatory Toxicology and Pharmacology 72, 292–309. 10.1016/j.yrtph.2015.04.010.

55. Alexander-Dann, B., Pruteanu, L.L., Oerton, E., Sharma, N., Berindan-Neagoe, I., Módos, D., and Bender, A. (2018). Developments in toxicogenomics: understanding and predicting compound-induced toxicity from gene expression data. Mol Omics 14, 218–236. 10.1039/C8MO00042E.

56. Harrill, J.A., Viant, M.R., Yauk, C.L., Sachana, M., Gant, T.W., Auerbach, S.S., Beger, R.D., Bouhifd, M., O’Brien, J., Burgoon, L., et al. (2021). Progress towards an OECD reporting framework for transcriptomics and metabolomics in regulatory toxicology. Regulatory Toxicology and Pharmacology 125, 105020. 10.1016/j.yrtph.2021.105020.

57. van Dam, S., Võsa, U., van der Graaf, A., Franke, L., and de Magalhães, J.P. (2018). Gene co-expression analysis for functional classification and gene-disease predictions. Brief Bioinform 19, 575–592. 10.1093/bib/bbw139.

58. Kumar, S. (2018). Cellular and molecular pathways of renal repair after acute kidney injury. Kidney Int 93, 27–40. 10.1016/j.kint.2017.07.030.

59. AbdulHameed, M.D.M., Ippolito, D.L., Stallings, J.D., and Wallqvist, A. (2016). Mining kidney toxicogenomic data by using gene co-expression modules. BMC Genomics 17, 790. 10.1186/s12864-016-3143-y.

60. Boulogne, F., Claus, L.R., Wiersma, H., Oelen, R., Schukking, F., de Klein, N., Li, S., Westra, H.-J., van der Zwaag, B., van Reekum, F., et al. (2023). KidneyNetwork: using kidney-derived gene expression data to predict and prioritize novel genes involved in kidney disease. European Journal of Human Genetics 31, 1300–1308. 10.1038/s41431-023-01296-x.

61. Jia, N.-Y., Liu, X.-Z., Zhang, Z., and Zhang, H. (2021). Weighted Gene Co-expression Network Analysis Reveals Different Immunity but Shared Renal Pathology Between IgA Nephropathy and Lupus Nephritis. Front Genet 12. 10.3389/fgene.2021.634171.

62. Xia, J., Hou, Y., Cai, A., Xu, Y., Yang, W., Huang, M., and Mou, S. (2023). An integrated co-expression network analysis reveals novel genetic biomarkers for immune cell infiltration in chronic kidney disease. Front Immunol 14. 10.3389/fimmu.2023.1129524.

63. Lin, X., Li, J., Tan, R., Zhong, X., Yang, J., and Wang, L. (2021). Identification of Hub Genes Associated with the Development of Acute Kidney Injury by Weighted Gene Co-Expression Network Analysis. Kidney Blood Press Res 46, 63–73. 10.1159/000511661.

64. Beckerman, P., Qiu, C., Park, J., Ledo, N., Ko, Y.-A., Park, A.-S.D., Han, S.-Y., Choi, P., Palmer, M., and Susztak, K. (2017). Human Kidney Tubule-Specific Gene Expression Based Dissection of Chronic Kidney Disease Traits. EBioMedicine 24, 267–276. 10.1016/j.ebiom.2017.09.014.

65. Arif, M., Zhang, C., Li, X., Güngör, C., Çakmak, B., Arslantürk, M., Tebani, A., Özcan, B., Subaş, O., Zhou, W., et al. (2021). iNetModels 2.0: an interactive visualization and database of multi-omics data. Nucleic Acids Res 49, W271–W276. 10.1093/nar/gkab254.

66. Raina, P., Guinea, R., Chatsirisupachai, K., Lopes, I., Farooq, Z., Guinea, C., Solyom, C.-A., and de Magalhães, J.P. (2023). GeneFriends: gene co-expression databases and tools for humans and model organisms. Nucleic Acids Res 51, D145–D158. 10.1093/nar/gkac1031.

67. Lee, S., Zhang, C., Arif, M., Liu, Z., Benfeitas, R., Bidkhori, G., Deshmukh, S., Al Shobky, M., Lovric, A., Boren, J., et al. (2018). TCSBN: a database of tissue and cancer specific biological networks. Nucleic Acids Res 46, D595–D600. 10.1093/nar/gkx994.

68. Lee, J., Shah, M., Ballouz, S., Crow, M., and Gillis, J. (2020). CoCoCoNet: conserved and comparative co-expression across a diverse set of species. Nucleic Acids Res 48, W566–W571. 10.1093/nar/gkaa348.

69. García-Ruiz, S., Gil-Martínez, A.L., Cisterna, A., Jurado-Ruiz, F., Reynolds, R.H., Cookson, M.R., Hardy, J., Ryten, M., and Botía, J.A. (2021). CoExp: A Web Tool for the Exploitation of Co-expression Networks. Front Genet 12. 10.3389/fgene.2021.630187.

70. Campos, G., Schmidt-Heck, W., De Smedt, J., Widera, A., Ghallab, A., Pütter, L., González, D., Edlund, K., Cadenas, C., Marchan, R., et al. (2020). Inflammation-associated suppression of metabolic gene networks in acute and chronic liver disease. Arch Toxicol 94, 205–217. 10.1007/s00204-019-02630-3.

71. Gudehithlu, K.P., Hart, P., Joshi, A., Garcia-Gomez, I., Cimbaluk, D.J., Dunea, G., Arruda, J.A.L., and Singh, A.K. (2019). Urine exosomal ceruloplasmin: a potential early biomarker of underlying kidney disease. Clin Exp Nephrol 23, 1013–1021. 10.1007/s10157-019-01734-5.

72. Hoffmann, D., Bijol, V., Krishnamoorthy, A., Gonzalez, V.R., Frendl, G., Zhang, Q., Goering, P.L., Brown, R.P., Waikar, S.S., and Vaidya, V.S. (2012). Fibrinogen Excretion in the Urine and Immunoreactivity in the Kidney Serves as a Translational Biomarker for Acute Kidney Injury. Am J Pathol 181, 818–828. 10.1016/j.ajpath.2012.06.004.

73. Arrizabalaga, P., Solé, M., Abellana, R., de las Cuevas, X., Soler, J., Pascual, J., and Ascaso, C. (2003). Tubular and Interstitial Expression of ICAM-1 as a Marker of Renal Injury in IgA Nephropathy. Am J Nephrol 23, 121–128. 10.1159/000068920.

74. Stern, E.P., Unwin, R., Burns, A., Ong, V.H., and Denton, C.P. (2021). Exploring molecular pathology of chronic kidney disease in systemic sclerosis by analysis of urinary and serum proteins. Rheumatol Adv Pract 5. 10.1093/rap/rkaa083.

75. Wei, J., Zhang, J., Wei, J., Hu, M., Chen, X., Qin, X., Chen, J., Lei, F., and Qin, Y. (2023). Identification of AGXT2, SHMT1, and ACO2 as important biomarkers of acute kidney injury by WGCNA. PLoS One 18, e0281439. 10.1371/journal.pone.0281439.

76. Wijaya, L.S., Kunnen, S.J., Trairatphisan, P., Fisher, C.P., Crosby, M.E., Schaefer, K., Bodié, K., Vaughan, E.E., Breidenbach, L., Reich, T., et al. (2025). Spatio-temporal transcriptomic analysis reveals distinct nephrotoxicity, DNA damage, and regeneration response after cisplatin. Cell Biol Toxicol 41, 49. 10.1007/s10565-025-10003-z.

77. Ferreira, S., Fisher, C., Furlong, L.I., Laplanche, L., Park, B.K., Pin, C., Saez-Rodriguez, J., and Trairatphisan, P. (2020). Quantitative Systems Toxicology Modeling To Address Key Safety Questions in Drug Development: A Focus of the TransQST Consortium. Chem Res Toxicol 33, 7–9. 10.1021/acs.chemrestox.9b00499.

