## Supplementary Figures for "Utilizing gene co-expression networks with the rat kidney TXG-MAPr tool to enhance safety assessment, biomarker identification and human translation"

**Figure S1. TXG-MAPr dendrograms.** TXG-MAPr dendrograms showing Ward's hierarchical clustering of the modules (nodes) in a linear (left) and circular (right) hierarchical tree. The dendrogram was cut in different branches (red dashed line) and the branch annotation is provided (*i.e.*, A1aIαi).

**Figure S2. Module TXG-MAPr dendrograms for cisplatin.** TXG-MAPr dendrograms showing WGCNA module eigengene scores (EGS) of the cisplatin treatment at all timepoints and dose levels. The size and colour of the circles is proportional to the module EGS, and the red/orange colours indicate induction of the module, while blue/green indicates repression of the module.

**Figure S3. Gene and module responses during nephrotoxic treatment conditions.** Module EGs (A) and gene log2 FC (B) of modules showing clear induction during nephrotoxic treatment conditions. (C) Module circle plot with the (top 25) gene members. The colour is proportional to the gene log2 FC for cisplatin treatment at 29 days, with red/orange colours indicating induction of the gene and blue/green indicating repression of the gene. APL = allopurinol, CSP = cisplatin, PAN = puromycin aminonucleoside. Key annotation of the modules is provided.

**Figure S4. Module association with pathology.** TXG-MAPr dendrogram indicating module association for different pathologies with a signed log10 adjusted p-value. Red colour indicates that the module EGS positively correlates with the selected pathology, while blue means a negative correlation (*i.e.,* module is repressed when the pathology is present).

**Figure S5. Heatmap of the module association with pathology.** Heatmap of signed log10 p-adjust of concurrent (A) and predictive (B) module associations with pathology. Red colour indicates that the module EGS positively correlates with the selected pathology, while blue means a negative correlation (*i.e.,* module is repressed when the pathology is present). Hierarchical clustering was applied on rows and columns to cluster both modules (rows) and toxicity phenotypes (columns), with similar association scores together. For concurrent associations, modules from clusters 1 and 2 were selected for strongest negative and positive correlation with toxicity, respectively. Module shows highest statistical association with concurrent toxicity phenotypes from cluster A, which were selected to look at the mean effect sizes and p-values (see also Table S9). For predictive associations, the 4 days show the highest statistical significance, where modules from clusters 2 and 7 were selected for strongest negative and positive correlation with toxicity, respectively. Modules in clusters 1 and 3 show the best association with 1-day predictive pathology. Heatmaps are clustered by Euclidean distance, using the complete method from *pheatmap* package.

**Figure S6. Statistics of the predictive module associations with pathology.** Box plots showing the median and quantiles of statistical parameters (effect size and log10 p-adjust) for predictive module association with pathology at different timepoints (x-axis). The predictive module association with toxicity showed on average higher effect sizes and p-values with increasing time. (A) Effect sizes are coloured for significant p-adjust values ($p < 0.05$). (B) Absolute log10 p-adjust values are coloured for significant effect sizes (Cohen's D > 0.8). Red line also indicates the significant effect sizes (A) and adjusted p-values (B), showing several modules with significant effect size and p-adjust (blue colour).

**Figure S7. Module TXG-MAPr dendrograms for nephrotoxic compounds.** TXG-MAPr dendrograms showing WGCNA module eigengene scores (EGS) of the nephrotoxic compounds at different timepoints and high dose levels. The size and colour of the circles is proportional to the module EGS, and the red/orange colours indicate induction of the module, while blue/green indicates repression of the module.

**Figure S8. Heatmap of module EGS for eight nephrotoxic compounds.** (A) Heatmap shows clustering of induced and repressed modules for the nephrotoxic drugs based on the module eigengene score. The colour is proportional to the module EGS, and the red/orange colours indicate induction of the module, while blue/green indicates repression of the module. (B) Heatmap on the bottom shows the histopathology scores (ranging from 0 = none to 4 = severe) for the different treatment conditions. Heatmap is clustered by Euclidean distance, using the complete method from *pheatmap* package.

**Figure S9. Treatment correlation of module EGS.** (A) Correlation plots of module EGS responses during cisplatin (CSP) and puromycin aminonucleoside (PAN) treatments at the same timepoints. Pearson R value is provided in the plot title. (B) Cluster correlation plot for nephrotoxic compounds shows strong Pearson R correlation between module EGS at time points when there is concurrent pathology (top left cluster in black box). Heatmap is clustered by Euclidean distance, using the complete method from *pheatmap* package. Left heatmap displays the percent increase of serum injury biomarkers (BUN and creatinine) compared to control (purple) and histopathology grades of the most prevalent kidney pathologies (blue), *e.g.,* necrosis, regeneration, dilatation, hyaline cast, cellular infiltration and fibrosis.

**Figure S10. Module EGS heatmaps of the DM data.** (A) Module EGS of all DM treatment conditions (columns) are shown in a heatmap and were clustered by Euclidean distance, using the Ward.D2 method from *pheatmap* package. Histopathology scores, BUN and serum creatinine levels are shown as column annotation for all treatment conditions, showing that nephrotoxic treatments cluster together on the left side of the heatmap. The pathology associations are shown as row annotation for each module. (B) Highlight of clusters 1, 2 and 5-9 of the heatmaps for the conditions with the most severe pathology to show clear deregulation and clustering of several modules.

**Figure S11. Correlation of ischemic reperfusion injury and primary PTC cultures.** (A) Correlation plots of module EGS during ischemic injury and nephrotoxic treatments cisplatin (CSP) and allopurinol (APL). (B) Correlation plots of module EGS during ischemic injury with valproic acid (VPA) or vehicle control (NaCl) pre-treatment. (C) Correlation plots of module EGS during PTC dedifferentiation and most similar nephrotoxic (APL, CSP and PAN) treatment conditions.

**Figure S12. Module responses of ischemic reperfusion injury and primary PTC cultures.** Heatmap of the strongest module EGS responses during ischemic injury and primary PTC dedifferentiation. Heatmap is clustered by Euclidean distance, using the complete method from *pheatmap* package.

**Figure S13. Heatmap of the modules EGS responses in human kidney samples.** Module EGS of human transplant samples (columns) are shown in a heatmap for the preserved modules (rows) in human with a EGS > 2 or EGS < -2. Heatmap is clustered by correlation distance, using the complete method from *pheatmap* package.

**Figure S14. Module and gene dynamics in human renal transplant samples.** Box plot for the different human transplant groups showing the module EGS (left) and the log2 fold change of the hub gene (second column) and most significant genes of each module. The groups are indicated by rejection type: non-rejecting, ABMR = antibody mediated rejection, TCMR = T-cell mediated rejection, MIXED = both antibody and T-cell mediated rejection. Significant differences between groups were calculated using a Wilcoxon test. Gene log2 FC are coloured by significance (blue colour with p-adjust < 0.05).

**Figure S1**

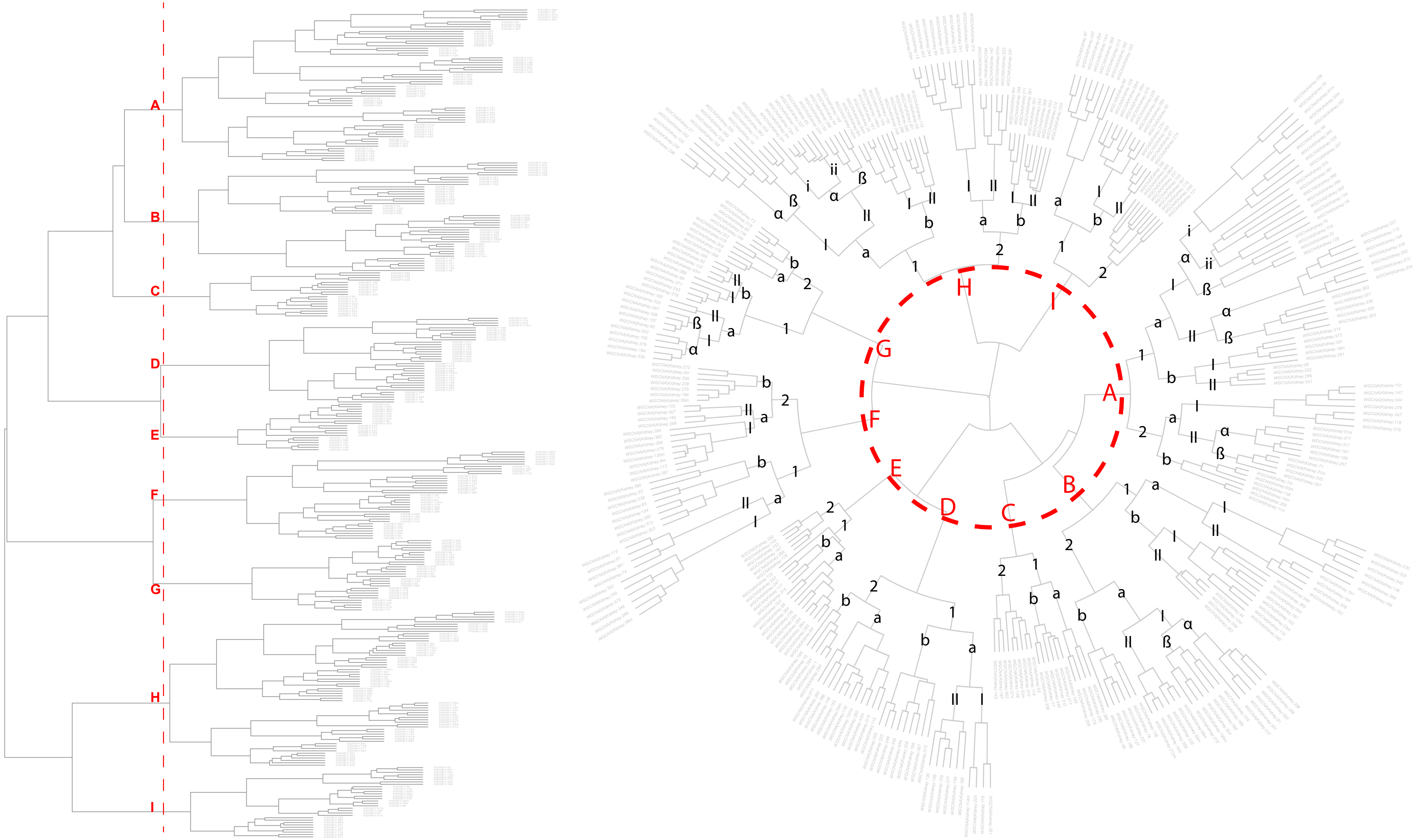

#### Figure S2

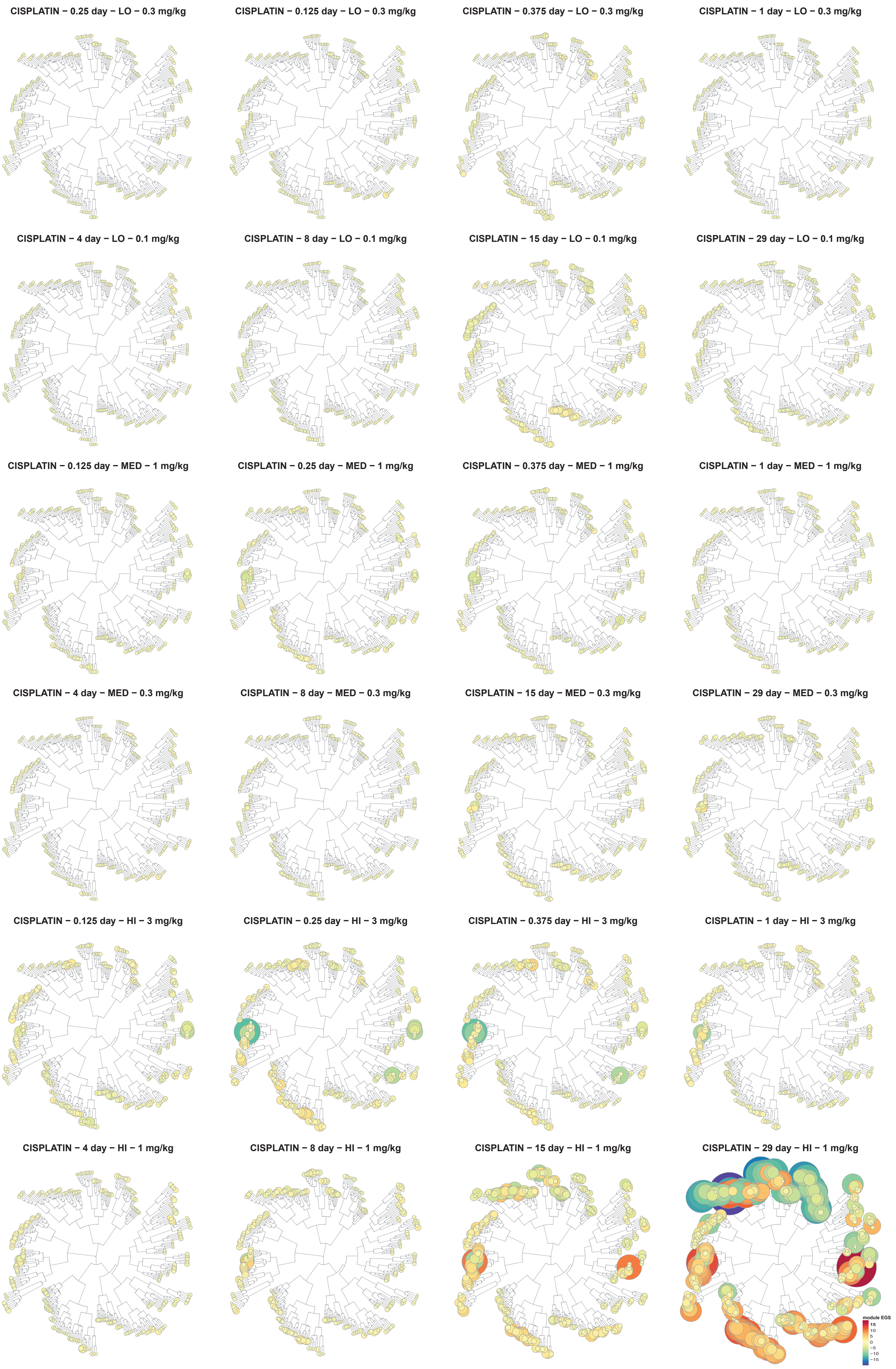
CISPLATIN − 0.25 day − LO − 0.3 mg/kg
CISPLATIN − 0.125 day − LO − 0.3 mg/kg
CISPLATIN − 0.375 day − LO − 0.3 mg/kg
CISPLATIN − 1 day − LO − 0.3 mg/kg
CISPLATIN − 4 day − LO − 0.1 mg/kg
CISPLATIN − 8 day − LO − 0.1 mg/kg
CISPLATIN − 15 day − LO − 0.1 mg/kg
CISPLATIN − 29 day − LO − 0.1 mg/kg
CISPLATIN − 0.125 day − MED − 1 mg/kg
CISPLATIN − 0.25 day − MED − 1 mg/kg
CISPLATIN − 0.375 day − MED − 1 mg/kg
CISPLATIN − 1 day − MED − 1 mg/kg
CISPLATIN − 4 day − MED − 0.3 mg/kg
CISPLATIN − 8 day − MED − 0.3 mg/kg
CISPLATIN − 15 day − MED − 0.3 mg/kg
CISPLATIN − 29 day − MED − 0.3 mg/kg
CISPLATIN − 0.125 day − HI − 3 mg/kg
CISPLATIN − 0.25 day − HI − 3 mg/kg
CISPLATIN − 0.375 day − HI − 3 mg/kg
CISPLATIN − 1 day − HI − 3 mg/kg
CISPLATIN − 4 day − HI − 1 mg/kg
CISPLATIN − 8 day − HI − 1 mg/kg
CISPLATIN − 15 day − HI − 1 mg/kg
CISPLATIN − 29 day − HI − 1 mg/kg
module EGS
15
10
5
0
−5
−10
−15

**Figure S3**

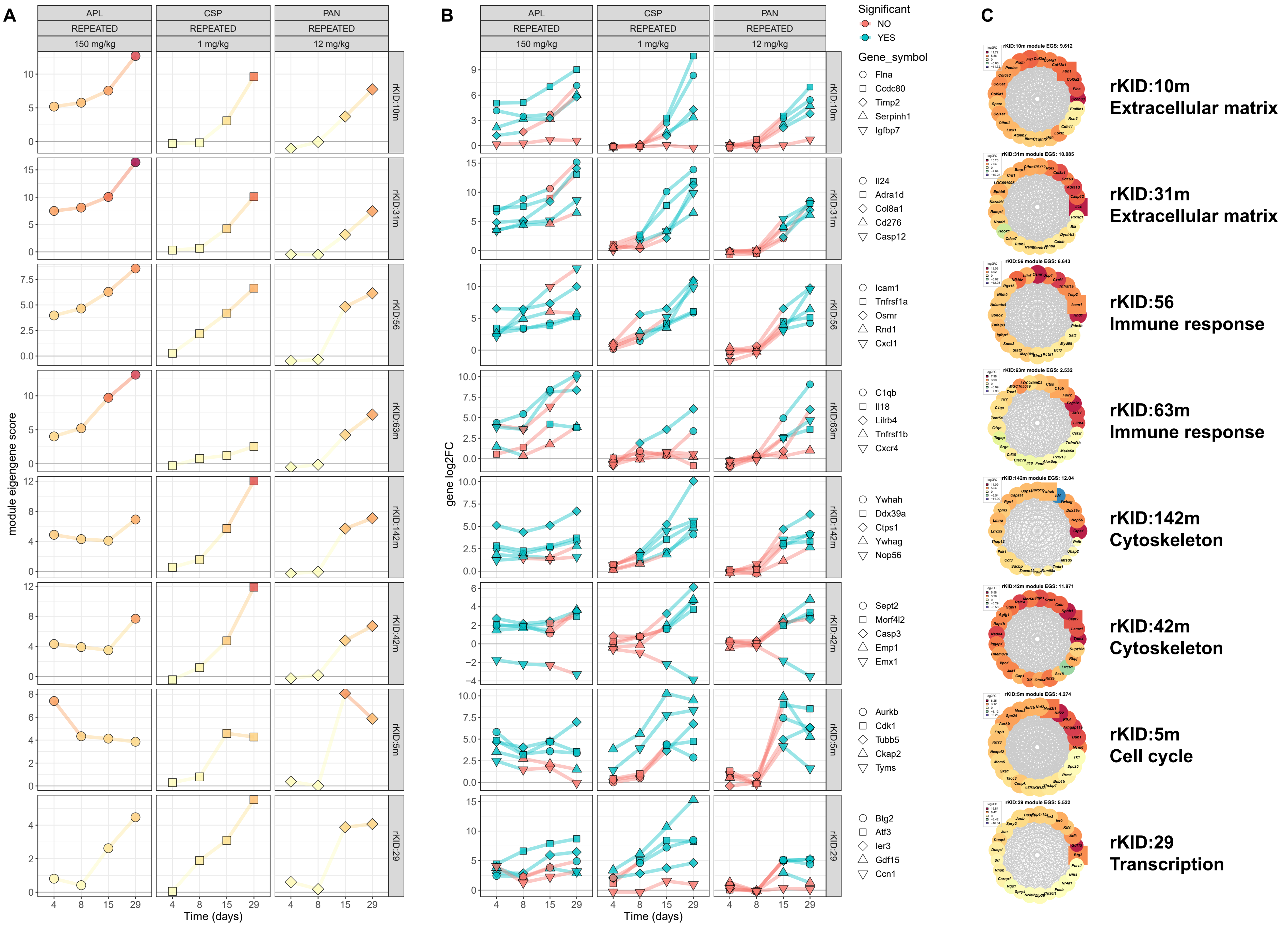

A
B
C
APL
CSP
PAN
REPEATED
150 mg/kg
1 mg/kg
12 mg/kg
rKID:10m
rKID:31m
rKID:56
rKID:63m
rKID:142m
rKID:42m
rKID:5m
rKID:29
module eigengene score
gene log2FC
Time (days)
Significant
NO
YES
Gene_symbol
Flna
Ccdc80
Timp2
Serpinh1
Igfbp7
Il24
Adra1d
Col8a1
Cd276
Casp12
Icam1
Tnfrsf1a
Osmr
Rnd1
Cxcl1
C1qb
Il18
Lilrb4
Tnfrsf1b
Cxcr4
Ywhah
Ddx39a
Ctps1
Ywhag
Nop56
Sept2
Morf4l2
Casp3
Emp1
Emx1
Aurkb
Cdk1
Tubb5
Ckap2
Tyms
Btg2
Atf3
Ier3
Gdf15
Ccn1
rKID:10m module EGS: 9.612
rKID:31m module EGS: 10.085
rKID:56 module EGS: 6.643
rKID:63m module EGS: 2.532
rKID:142m module EGS: 12.04
rKID:42m module EGS: 11.871
rKID:5m module EGS: 4.274
rKID:29 module EGS: 5.522
rKID:10m
Extracellular matrix
rKID:31m
Extracellular matrix
rKID:56
Immune response
rKID:63m
Immune response
rKID:142m
Cytoskeleton
rKID:42m
Cytoskeleton
rKID:5m
Cell cycle
rKID:29
Transcription

### Figure S4

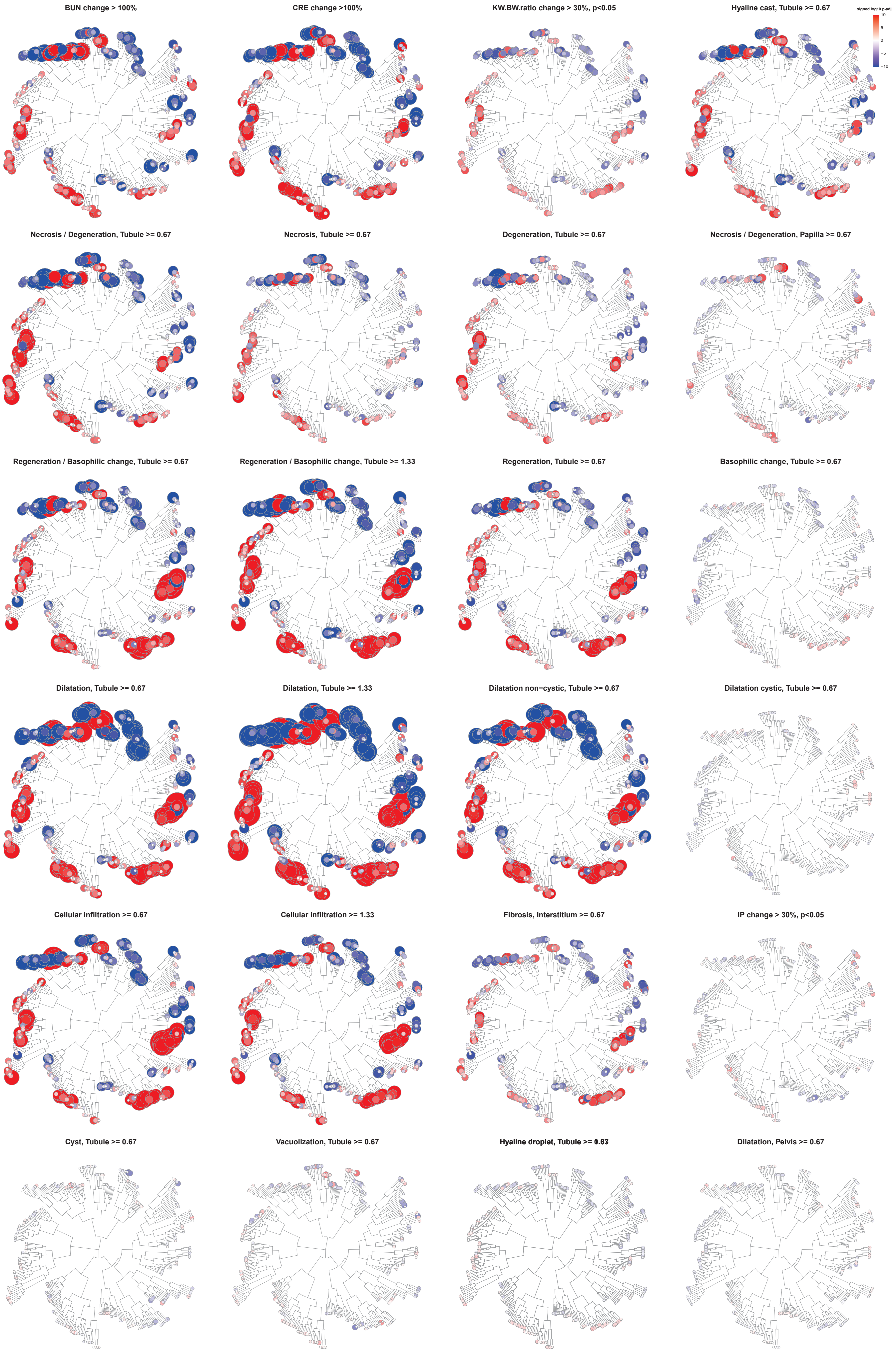
BUN change > 100%
CRE change >100%
KW.BW.ratio change > 30%, p<0.05
Hyaline cast, Tubule >= 0.67
signed log10 p-adj
10
5
0
−5
−10
Necrosis / Degeneration, Tubule >= 0.67
Necrosis, Tubule >= 0.67
Degeneration, Tubule >= 0.67
Necrosis / Degeneration, Papilla >= 0.67
Regeneration / Basophilic change, Tubule >= 0.67
Regeneration / Basophilic change, Tubule >= 1.33
Regeneration, Tubule >= 0.67
Basophilic change, Tubule >= 0.67
Dilatation, Tubule >= 0.67
Dilatation, Tubule >= 1.33
Dilatation non−cystic, Tubule >= 0.67
Dilatation cystic, Tubule >= 0.67
Cellular infiltration >= 0.67
Cellular infiltration >= 1.33
Fibrosis, Interstitium >= 0.67
IP change > 30%, p<0.05
Cyst, Tubule >= 0.67
Vacuolization, Tubule >= 0.67
Hyaline droplet, Tubule >= 0.67
Dilatation, Pelvis >= 0.67

**Figure S5**

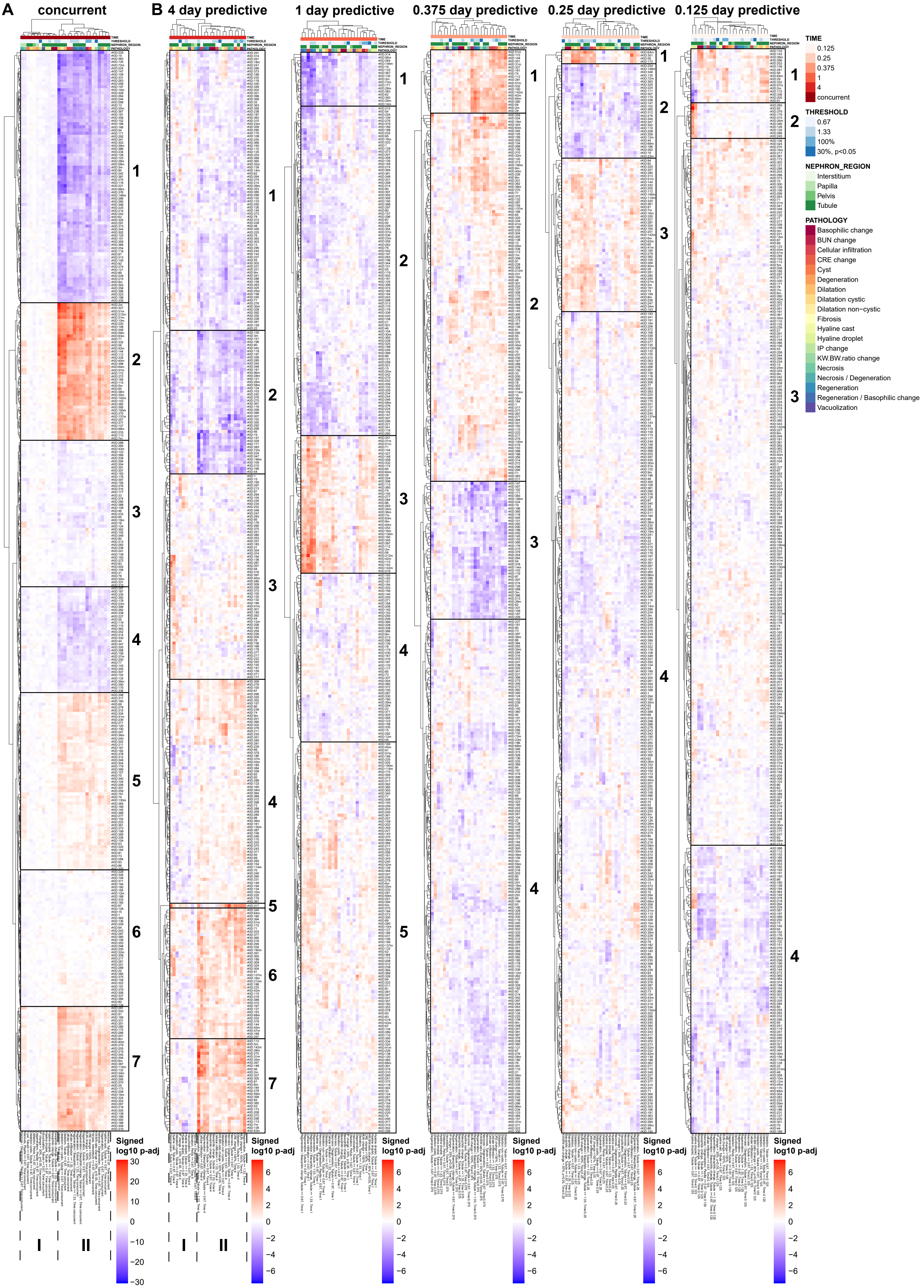
A
concurrent
B
4 day predictive
1 day predictive
0.375 day predictive
0.25 day predictive
0.125 day predictive
TIME
0.125
0.25
0.375
1
4
concurrent
THRESHOLD
0.67
1.33
100%
30%, p<0.05
NEPHRON_REGION
Interstitium
Papilla
Pelvis
Tubule
PATHOLOGY
Basophilic change
BUN change
Cellular infiltration
CRE change
Cyst
Degeneration
Dilatation
Dilatation cystic
Dilatation non−cystic
Fibrosis
Hyaline cast
Hyaline droplet
IP change
KW.BW.ratio change
Necrosis
Necrosis / Degeneration
Regeneration
Regeneration / Basophilic change
Vacuolization
Signed
log10 p-adj
I
II

#### Figure S6

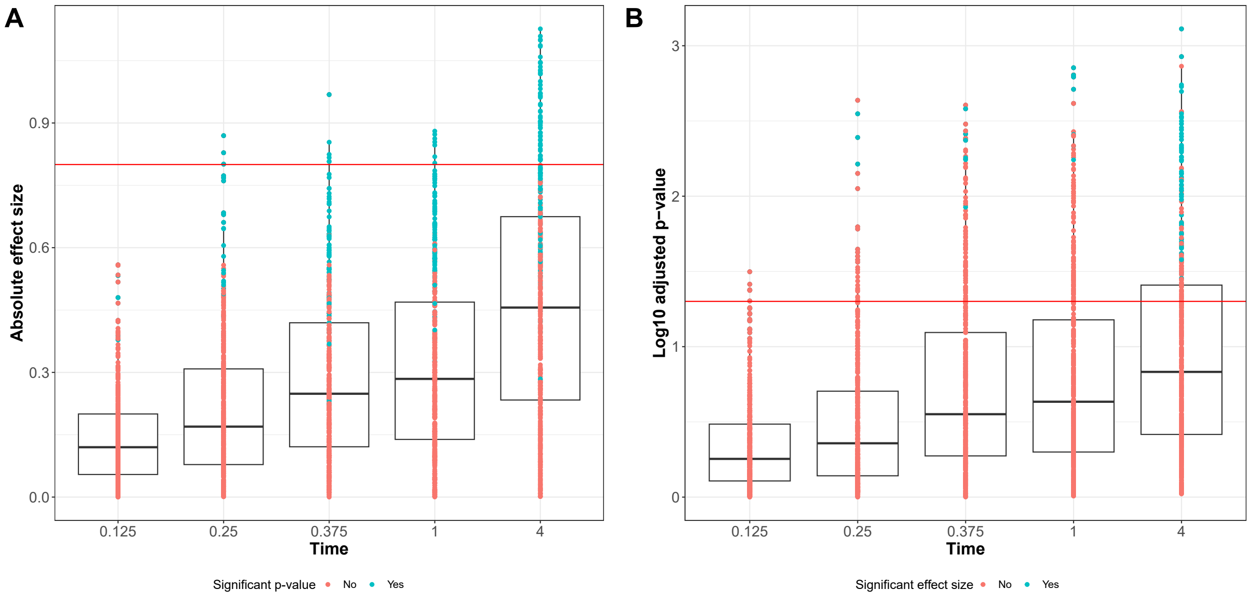
A
Absolute effect size
0.0
0.3
0.6
0.9
0.125
0.25
0.375
1
4
Time
Significant p-value
No
Yes
B
Log10 adjusted p−value
0
1
2
3
0.125
0.25
0.375
1
4
Time
Significant effect size
No
Yes

#### Figure S7

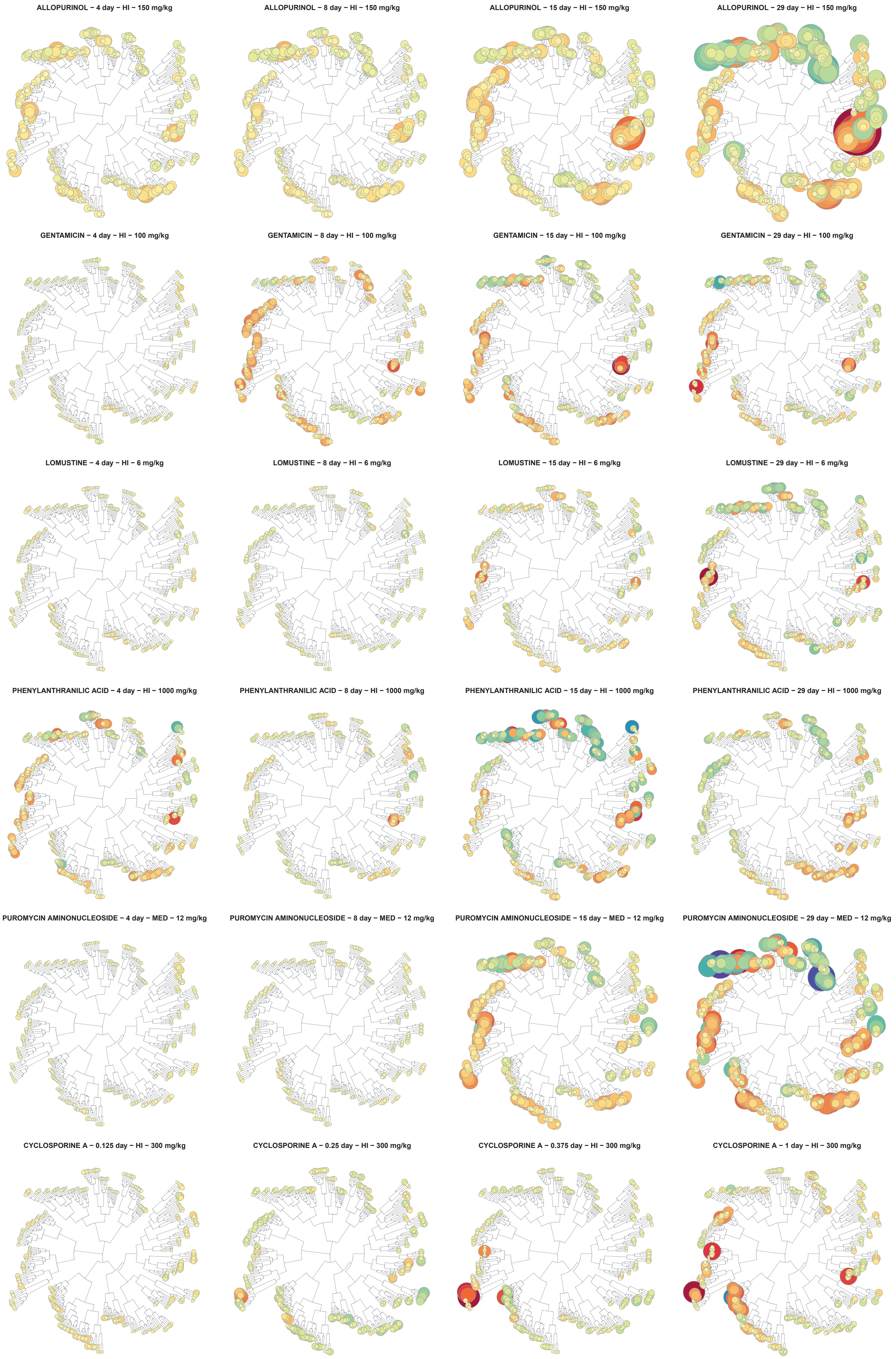

ALLOPURINOL – 4 day – HI – 150 mg/kg
ALLOPURINOL – 8 day – HI – 150 mg/kg
ALLOPURINOL – 15 day – HI – 150 mg/kg
ALLOPURINOL – 29 day – HI – 150 mg/kg
GENTAMICIN – 4 day – HI – 100 mg/kg
GENTAMICIN – 8 day – HI – 100 mg/kg
GENTAMICIN – 15 day – HI – 100 mg/kg
GENTAMICIN – 29 day – HI – 100 mg/kg
LOMUSTINE – 4 day – HI – 6 mg/kg
LOMUSTINE – 8 day – HI – 6 mg/kg
LOMUSTINE – 15 day – HI – 6 mg/kg
LOMUSTINE – 29 day – HI – 6 mg/kg
PHENYLANTHRANILIC ACID – 4 day – HI – 1000 mg/kg
PHENYLANTHRANILIC ACID – 8 day – HI – 1000 mg/kg
PHENYLANTHRANILIC ACID – 15 day – HI – 1000 mg/kg
PHENYLANTHRANILIC ACID – 29 day – HI – 1000 mg/kg
PUROMYCIN AMINONUCLEOSIDE – 4 day – MED – 12 mg/kg
PUROMYCIN AMINONUCLEOSIDE – 8 day – MED – 12 mg/kg
PUROMYCIN AMINONUCLEOSIDE – 15 day – MED – 12 mg/kg
PUROMYCIN AMINONUCLEOSIDE – 29 day – MED – 12 mg/kg
CYCLOSPORINE A – 0.125 day – HI – 300 mg/kg
CYCLOSPORINE A – 0.25 day – HI – 300 mg/kg
CYCLOSPORINE A – 0.375 day – HI – 300 mg/kg
CYCLOSPORINE A – 1 day – HI – 300 mg/kg

### Figure S8

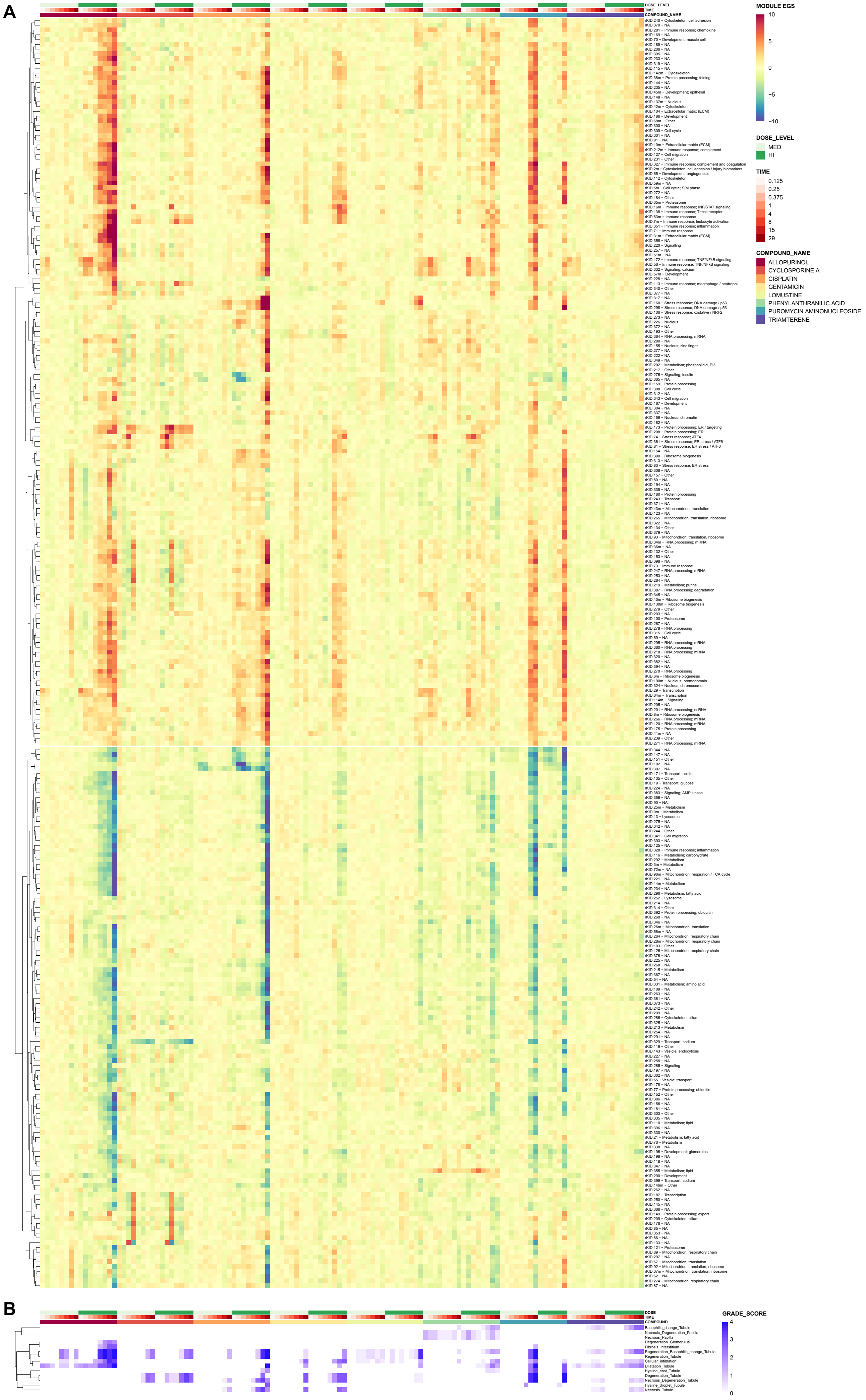
A
B
DOSE_LEVEL
TIME
COMPOUND_NAME
MODULE EGS
10
5
0
−5
−10
DOSE_LEVEL
MED
HI
TIME
0.125
0.25
0.375
1
4
8
15
29
COMPOUND_NAME
ALLOPURINOL
CYCLOSPORINE A
CISPLATIN
GENTAMICIN
LOMUSTINE
PHENYLANTHRANILIC ACID
PUROMYCIN AMINONUCLEOSIDE
TRIAMTERENE
DOSE
TIME
COMPOUND
Basophilic_change_Tubule
Necrosis_Degeneration_Papilla
Necrosis_Papilla
Degeneration_Glomerulus
Fibrosis_Interstitium
Regeneration_Basophilic_change_Tubule
Regeneration_Tubule
Cellular_infiltration
Dilatation_Tubule
Hyaline_cast_Tubule
Degeneration_Tubule
Necrosis_Degeneration_Tubule
Hyaline_droplet_Tubule
Necrosis_Tubule
GRADE_SCORE
4
3
2
1
0

#### Figure S9

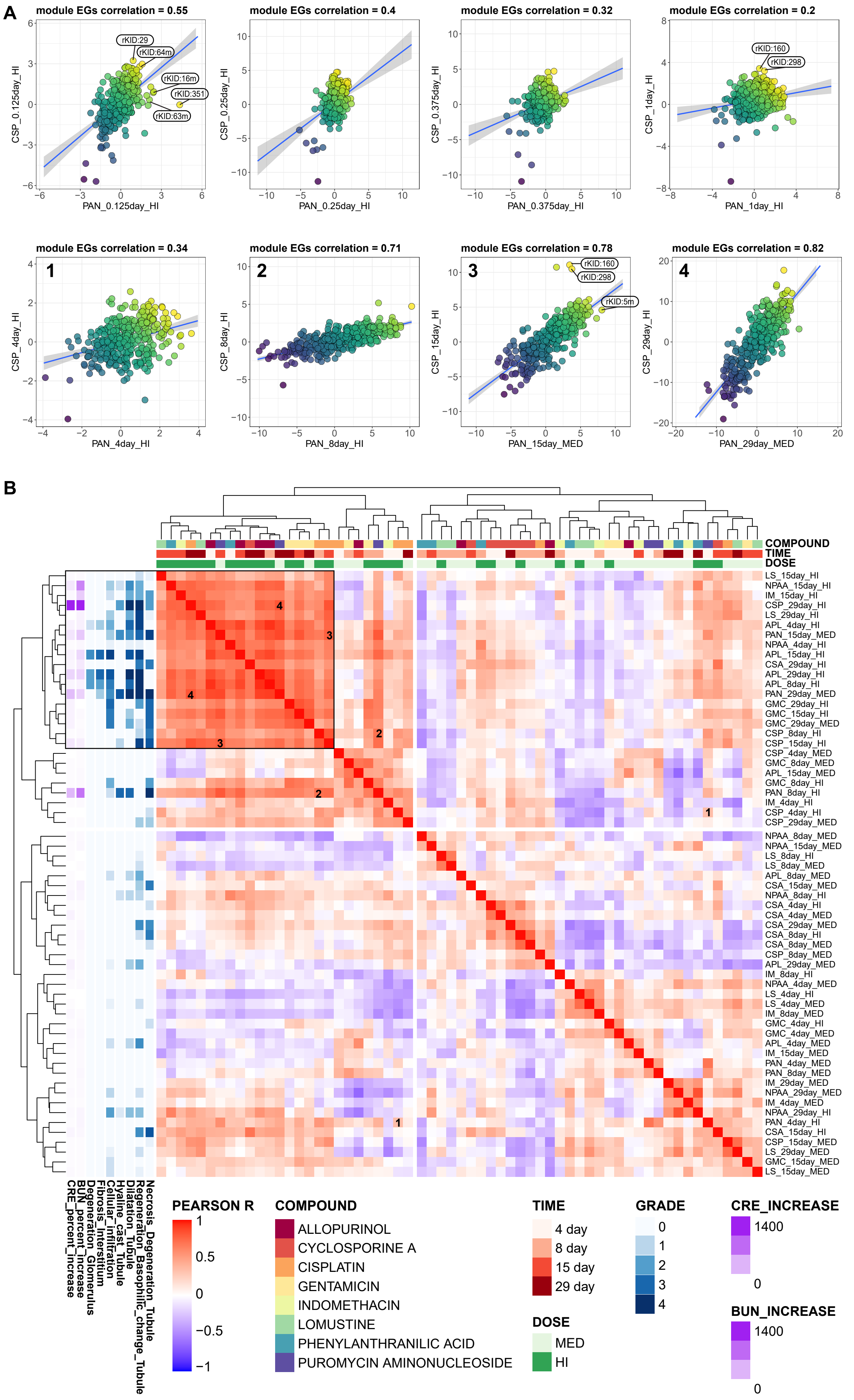

A
module EGs correlation = 0.55
CSP_0.125day_HI
PAN_0.125day_HI
rKID:29
rKID:64m
rKID:16m
rKID:351
rKID:63m
module EGs correlation = 0.4
CSP_0.25day_HI
PAN_0.25day_HI
module EGs correlation = 0.32
CSP_0.375day_HI
PAN_0.375day_HI
module EGs correlation = 0.2
CSP_1day_HI
PAN_1day_HI
rKID:160
rKID:298
module EGs correlation = 0.34
CSP_4day_HI
PAN_4day_HI
module EGs correlation = 0.71
CSP_8day_HI
PAN_8day_HI
module EGs correlation = 0.78
CSP_15day_HI
PAN_15day_MED
rKID:160
rKID:298
rKID:5m
module EGs correlation = 0.82
CSP_29day_HI
PAN_29day_MED
B
COMPOUND
TIME
DOSE
LS_15day_HI
NPAA_15day_HI
IM_15day_HI
CSP_29day_HI
LS_29day_HI
APL_4day_HI
PAN_15day_MED
NPAA_4day_HI
APL_15day_HI
CSA_29day_HI
APL_29day_HI
APL_8day_HI
PAN_29day_MED
GMC_29day_HI
GMC_15day_HI
GMC_29day_MED
CSP_8day_HI
CSP_15day_HI
CSP_4day_MED
GMC_8day_MED
APL_15day_MED
GMC_8day_HI
PAN_8day_HI
IM_4day_HI
CSP_4day_HI
CSP_29day_MED
NPAA_8day_MED
NPAA_15day_MED
LS_8day_HI
LS_8day_MED
APL_8day_MED
CSA_15day_MED
NPAA_8day_HI
CSA_4day_HI
CSA_4day_MED
CSA_29day_MED
CSA_8day_HI
CSA_8day_MED
CSP_8day_MED
APL_29day_MED
IM_8day_HI
NPAA_4day_MED
LS_4day_HI
LS_4day_MED
IM_8day_MED
GMC_4day_HI
GMC_4day_MED
APL_4day_MED
IM_15day_MED
PAN_4day_MED
PAN_8day_MED
IM_29day_MED
NPAA_29day_MED
IM_4day_MED
NPAA_29day_HI
PAN_4day_HI
CSA_15day_HI
CSP_15day_MED
LS_29day_MED
GMC_15day_MED
LS_15day_MED
Necrosis_Degeneration_Tubule
Regeneration_Basophilic_change_Tubule
Dilatation_Tubule
Hyaline_cast_Tubule
Cellular_infiltration
Fibrosis_Interstitium
Degeneration_Glomerulus
BUN_percent_increase
CRE_percent_increase
PEARSON R
COMPOUND
ALLOPURINOL
CYCLOSPORINE A
CISPLATIN
GENTAMICIN
INDOMETHACIN
LOMUSTINE
PHENYLANTHRANILIC ACID
PUROMYCIN AMINONUCLEOSIDE
TIME
4 day
8 day
15 day
29 day
DOSE
MED
HI
GRADE
CRE_INCREASE
1400
BUN_INCREASE
1400

#### Figure S10

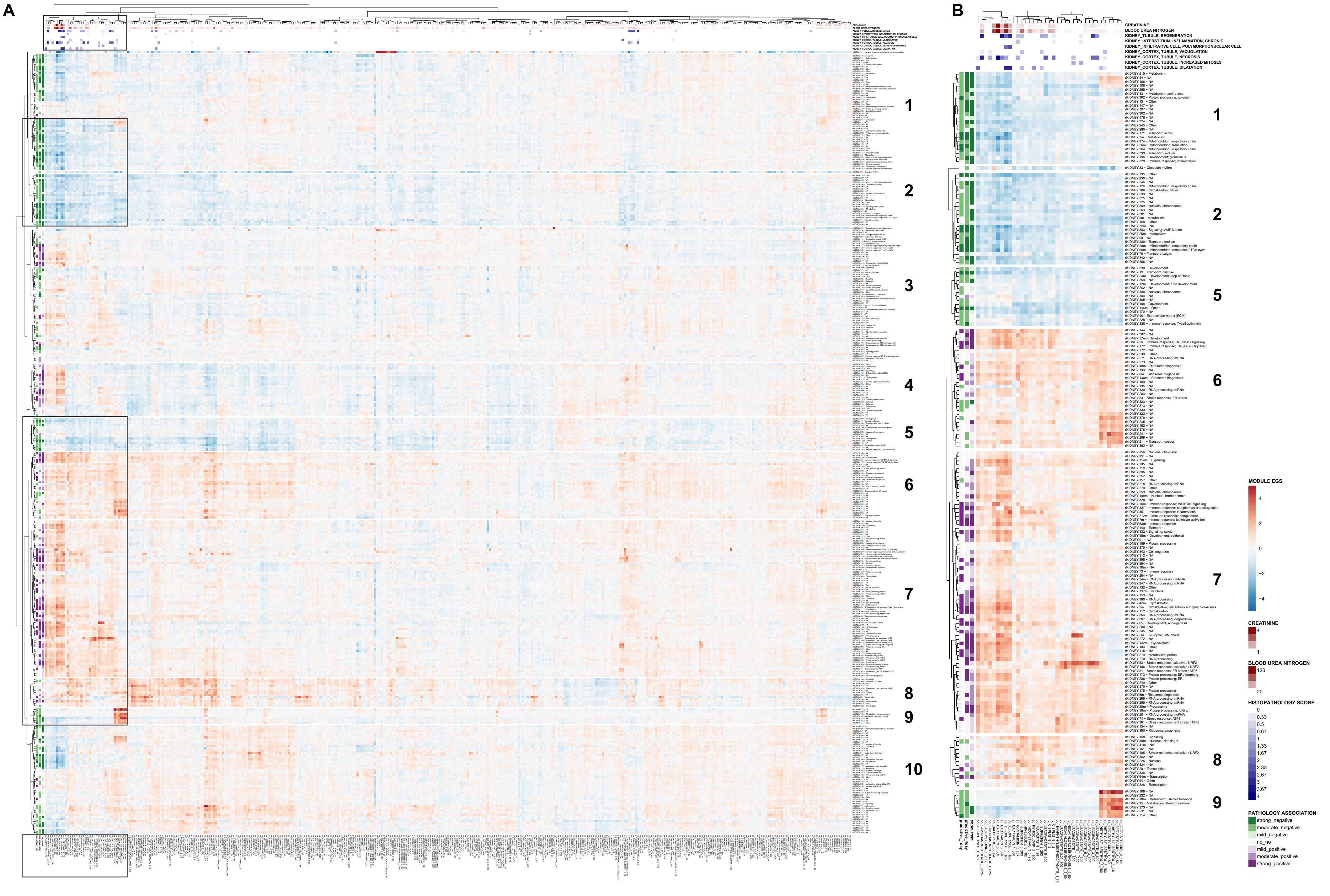
A
B
CREATININE
BLOOD UREA NITROGEN
KIDNEY_TUBULE, REGENERATION
KIDNEY_INTERSTITIUM, INFLAMMATION, CHRONIC
KIDNEY_INFILTRATIVE CELL, POLYMORPHONUCLEAR CELL
KIDNEY_CORTEX, TUBULE, VACUOLATION
KIDNEY_CORTEX, TUBULE, NECROSIS
KIDNEY_CORTEX, TUBULE, INCREASED MITOSES
KIDNEY_CORTEX, TUBULE, DILATATION
1
2
3
4
5
6
7
8
9
10
concurrent
predictive_4day
predictive_1day
MODULE EGS
4
2
0
-2
-4
CREATININE
4
1
BLOOD UREA NITROGEN
120
20
HISTOPATHOLOGY SCORE
0
0.33
0.5
0.67
1
1.33
1.67
2
2.33
2.67
3
3.67
4
PATHOLOGY ASSOCIATION
strong_negative
moderate_negative
mild_negative
no_no
mild_positive
moderate_positive
strong_positive

#### Figure S11

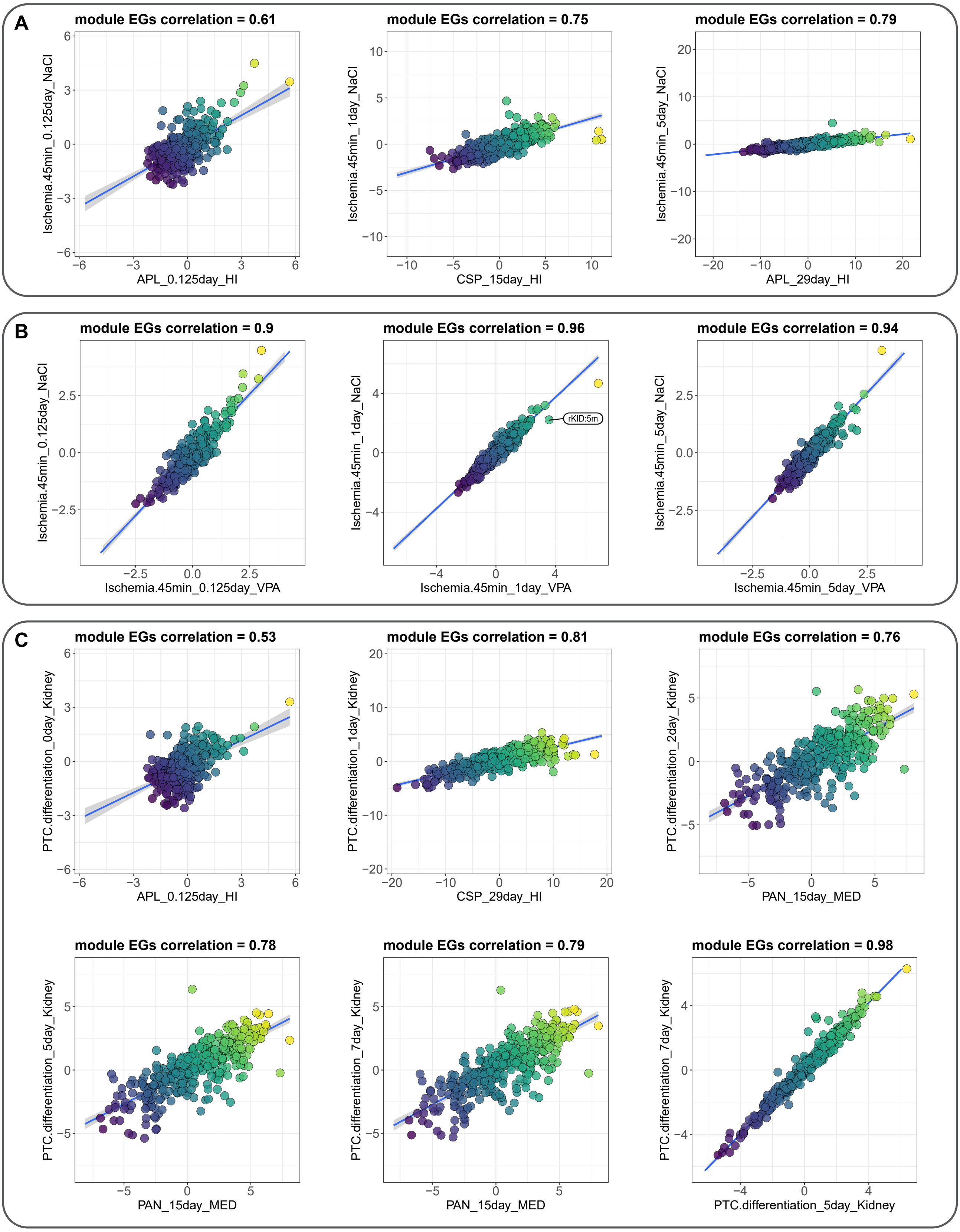
A
module EGs correlation = 0.61
Ischemia.45min_0.125day_NaCl
APL_0.125day_HI
module EGs correlation = 0.75
Ischemia.45min_1day_NaCl
CSP_15day_HI
module EGs correlation = 0.79
Ischemia.45min_5day_NaCl
APL_29day_HI
B
module EGs correlation = 0.9
Ischemia.45min_0.125day_NaCl
Ischemia.45min_0.125day_VPA
module EGs correlation = 0.96
Ischemia.45min_1day_NaCl
Ischemia.45min_1day_VPA
rKID:5m
module EGs correlation = 0.94
Ischemia.45min_5day_NaCl
Ischemia.45min_5day_VPA
C
module EGs correlation = 0.53
PTC.differentiation_0day_Kidney
APL_0.125day_HI
module EGs correlation = 0.81
PTC.differentiation_1day_Kidney
CSP_29day_HI
module EGs correlation = 0.76
PTC.differentiation_2day_Kidney
PAN_15day_MED
module EGs correlation = 0.78
PTC.differentiation_5day_Kidney
PAN_15day_MED
module EGs correlation = 0.79
PTC.differentiation_7day_Kidney
PAN_15day_MED
module EGs correlation = 0.98
PTC.differentiation_7day_Kidney
PTC.differentiation_5day_Kidney

**Figure S12**

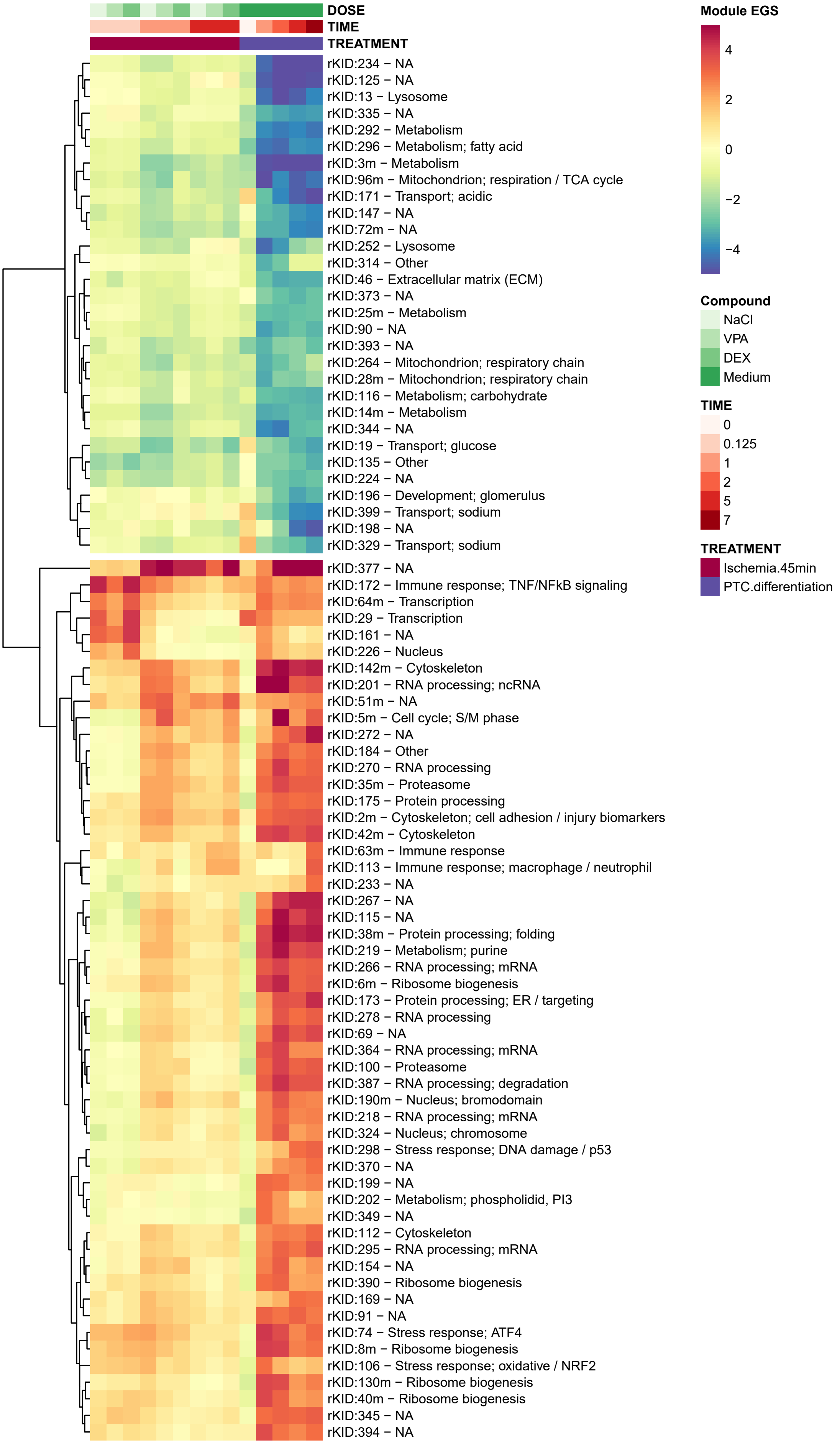
DOSE
TIME
TREATMENT
rKID:234 − NA
rKID:125 − NA
rKID:13 − Lysosome
rKID:335 − NA
rKID:292 − Metabolism
rKID:296 − Metabolism; fatty acid
rKID:3m − Metabolism
rKID:96m − Mitochondrion; respiration / TCA cycle
rKID:171 − Transport; acidic
rKID:147 − NA
rKID:72m − NA
rKID:252 − Lysosome
rKID:314 − Other
rKID:46 − Extracellular matrix (ECM)
rKID:373 − NA
rKID:25m − Metabolism
rKID:90 − NA
rKID:393 − NA
rKID:264 − Mitochondrion; respiratory chain
rKID:28m − Mitochondrion; respiratory chain
rKID:116 − Metabolism; carbohydrate
rKID:14m − Metabolism
rKID:344 − NA
rKID:19 − Transport; glucose
rKID:135 − Other
rKID:224 − NA
rKID:196 − Development; glomerulus
rKID:399 − Transport; sodium
rKID:198 − NA
rKID:329 − Transport; sodium
rKID:377 − NA
rKID:172 − Immune response; TNF/NFkB signaling
rKID:64m − Transcription
rKID:29 − Transcription
rKID:161 − NA
rKID:226 − Nucleus
rKID:142m − Cytoskeleton
rKID:201 − RNA processing; ncRNA
rKID:51m − NA
rKID:5m − Cell cycle; S/M phase
rKID:272 − NA
rKID:184 − Other
rKID:270 − RNA processing
rKID:35m − Proteasome
rKID:175 − Protein processing
rKID:2m − Cytoskeleton; cell adhesion / injury biomarkers
rKID:42m − Cytoskeleton
rKID:63m − Immune response
rKID:113 − Immune response; macrophage / neutrophil
rKID:233 − NA
rKID:267 − NA
rKID:115 − NA
rKID:38m − Protein processing; folding
rKID:219 − Metabolism; purine
rKID:266 − RNA processing; mRNA
rKID:6m − Ribosome biogenesis
rKID:173 − Protein processing; ER / targeting
rKID:278 − RNA processing
rKID:69 − NA
rKID:364 − RNA processing; mRNA
rKID:100 − Proteasome
rKID:387 − RNA processing; degradation
rKID:190m − Nucleus; bromodomain
rKID:218 − RNA processing; mRNA
rKID:324 − Nucleus; chromosome
rKID:298 − Stress response; DNA damage / p53
rKID:370 − NA
rKID:199 − NA
rKID:202 − Metabolism; phospholidid, PI3
rKID:349 − NA
rKID:112 − Cytoskeleton
rKID:295 − RNA processing; mRNA
rKID:154 − NA
rKID:390 − Ribosome biogenesis
rKID:169 − NA
rKID:91 − NA
rKID:74 − Stress response; ATF4
rKID:8m − Ribosome biogenesis
rKID:106 − Stress response; oxidative / NRF2
rKID:130m − Ribosome biogenesis
rKID:40m − Ribosome biogenesis
rKID:345 − NA
rKID:394 − NA
Module EGS
4
2
0
−2
−4
Compound
NaCl
VPA
DEX
Medium
TIME
0
0.125
1
2
5
7
TREATMENT
Ischemia.45min
PTC.differentiation

### Figure S13

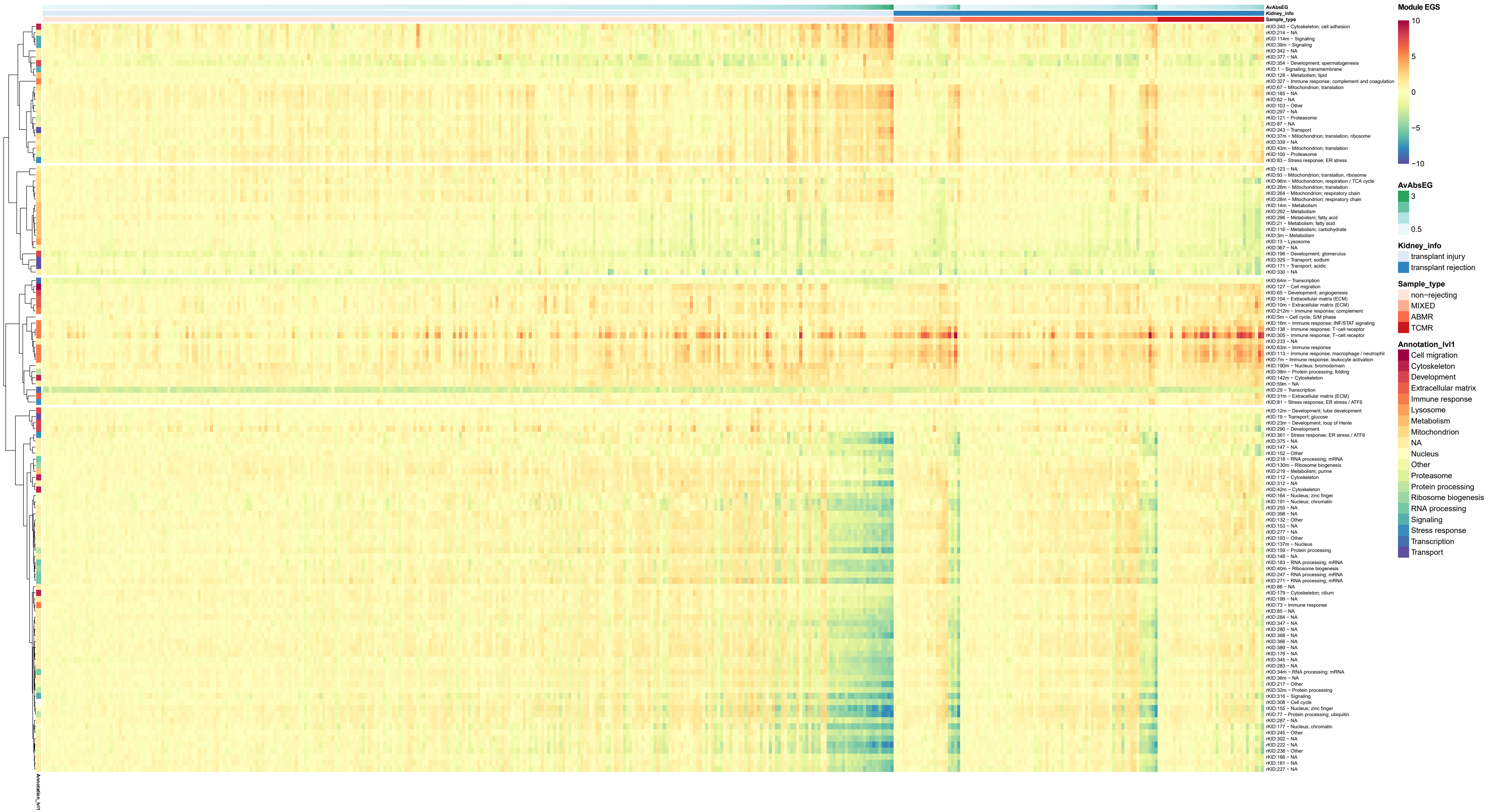
AvAbsEG
Kidney_info
Sample_type
rKID:240 − Cytoskeleton; cell adhesion
rKID:214 − NA
rKID:114m − Signaling
rKID:39m − Signaling
rKID:342 − NA
rKID:377 − NA
rKID:354 − Development; spermatogenesis
rKID:1 − Signaling; transmembrane
rKID:128 − Metabolism; lipid
rKID:327 − Immune response; complement and coagulation
rKID:67 − Mitochondrion; translation
rKID:185 − NA
rKID:62 − NA
rKID:103 − Other
rKID:297 − NA
rKID:121 − Proteasome
rKID:87 − NA
rKID:243 − Transport
rKID:37m − Mitochondrion; translation, ribosome
rKID:339 − NA
rKID:43m − Mitochondrion; translation
rKID:100 − Proteasome
rKID:83 − Stress response; ER stress
rKID:123 − NA
rKID:93 − Mitochondrion; translation, ribosome
rKID:96m − Mitochondrion; respiration / TCA cycle
rKID:26m − Mitochondrion; translation
rKID:264 − Mitochondrion; respiratory chain
rKID:28m − Mitochondrion; respiratory chain
rKID:14m − Metabolism
rKID:292 − Metabolism
rKID:296 − Metabolism; fatty acid
rKID:21 − Metabolism; fatty acid
rKID:116 − Metabolism; carbohydrate
rKID:3m − Metabolism
rKID:13 − Lysosome
rKID:367 − NA
rKID:196 − Development; glomerulus
rKID:329 − Transport; sodium
rKID:171 − Transport; acidic
rKID:330 − NA
rKID:64m − Transcription
rKID:127 − Cell migration
rKID:65 − Development; angiogenesis
rKID:104 − Extracellular matrix (ECM)
rKID:10m − Extracellular matrix (ECM)
rKID:212m − Immune response; complement
rKID:5m − Cell cycle; S/M phase
rKID:16m − Immune response; INF/STAT signaling
rKID:138 − Immune response; T−cell receptor
rKID:305 − Immune response; T−cell receptor
rKID:233 − NA
rKID:63m − Immune response
rKID:113 − Immune response; macrophage / neutrophil
rKID:7m − Immune response; leukocyte activation
rKID:190m − Nucleus; bromodomain
rKID:38m − Protein processing; folding
rKID:142m − Cytoskeleton
rKID:59m − NA
rKID:29 − Transcription
rKID:31m − Extracellular matrix (ECM)
rKID:81 − Stress response; ER stress / ATF6
rKID:12m − Development; tube development
rKID:19 − Transport; glucose
rKID:23m − Development; loop of Henle
rKID:290 − Development
rKID:361 − Stress response; ER stress / ATF6
rKID:375 − NA
rKID:147 − NA
rKID:152 − Other
rKID:218 − RNA processing; mRNA
rKID:130m − Ribosome biogenesis
rKID:219 − Metabolism; purine
rKID:112 − Cytoskeleton
rKID:312 − NA
rKID:42m − Cytoskeleton
rKID:164 − Nucleus; zinc finger
rKID:191 − Nucleus; chromatin
rKID:255 − NA
rKID:398 − NA
rKID:132 − Other
rKID:153 − NA
rKID:277 − NA
rKID:193 − Other
rKID:137m − Nucleus
rKID:159 − Protein processing
rKID:148 − NA
rKID:183 − RNA processing; mRNA
rKID:40m − Ribosome biogenesis
rKID:247 − RNA processing; mRNA
rKID:271 − RNA processing; mRNA
rKID:86 − NA
rKID:179 − Cytoskeleton; cilium
rKID:199 − NA
rKID:73 − Immune response
rKID:85 − NA
rKID:284 − NA
rKID:347 − NA
rKID:280 − NA
rKID:368 − NA
rKID:366 − NA
rKID:389 − NA
rKID:176 − NA
rKID:345 − NA
rKID:283 − NA
rKID:34m − RNA processing; mRNA
rKID:36m − NA
rKID:217 − Other
rKID:32m − Protein processing
rKID:316 − Signaling
rKID:308 − Cell cycle
rKID:155 − Nucleus; zinc finger
rKID:77 − Protein processing; ubiquitin
rKID:287 − NA
rKID:177 − Nucleus; chromatin
rKID:245 − Other
rKID:302 − NA
rKID:222 − NA
rKID:238 − Other
rKID:166 − NA
rKID:181 − NA
rKID:227 − NA
Annotation_lvl1
Module EGS
10
5
0
−5
−10
AvAbsEG
3
0.5
Kidney_info
transplant injury
transplant rejection
Sample_type
non−rejecting
MIXED
ABMR
TCMR
Annotation_lvl1
Cell migration
Cytoskeleton
Development
Extracellular matrix
Immune response
Lysosome
Metabolism
Mitochondrion
NA
Nucleus
Other
Proteasome
Protein processing
Ribosome biogenesis
RNA processing
Signaling
Stress response
Transcription
Transport

### Figure S14

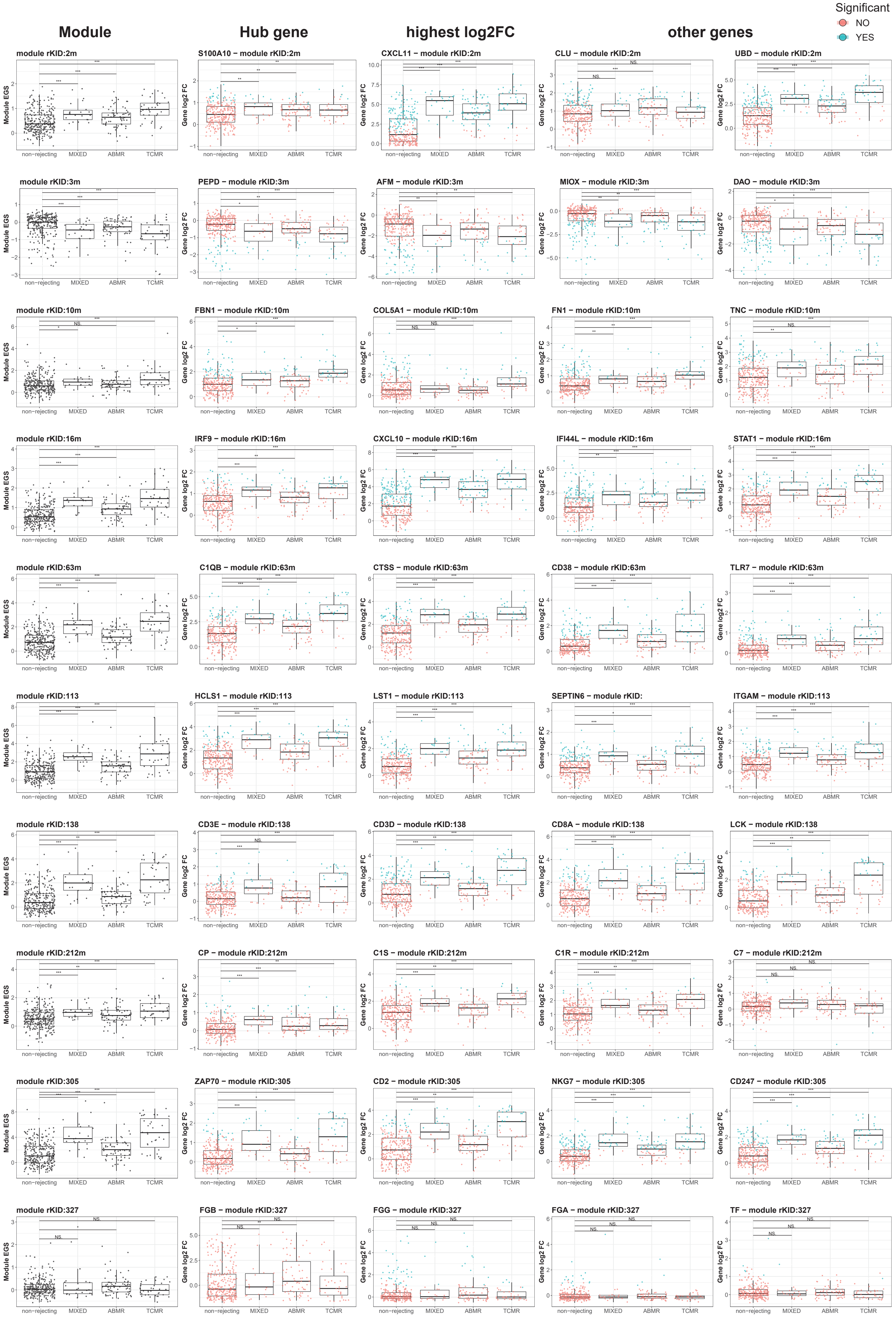
Module
Hub gene
highest log2FC
other genes
Significant
NO
YES
module rKID:2m
S100A10 – module rKID:2m
CXCL11 – module rKID:2m
CLU – module rKID:2m
UBD – module rKID:2m
module rKID:3m
PEPD – module rKID:3m
AFM – module rKID:3m
MIOX – module rKID:3m
DAO – module rKID:3m
module rKID:10m
FBN1 – module rKID:10m
COL5A1 – module rKID:10m
FN1 – module rKID:10m
TNC – module rKID:10m
module rKID:16m
IRF9 – module rKID:16m
CXCL10 – module rKID:16m
IFI44L – module rKID:16m
STAT1 – module rKID:16m
module rKID:63m
C1QB – module rKID:63m
CTSS – module rKID:63m
CD38 – module rKID:63m
TLR7 – module rKID:63m
module rKID:113
HCLS1 – module rKID:113
LST1 – module rKID:113
SEPTIN6 – module rKID:
ITGAM – module rKID:113
module rKID:138
CD3E – module rKID:138
CD3D – module rKID:138
CD8A – module rKID:138
LCK – module rKID:138
module rKID:212m
CP – module rKID:212m
C1S – module rKID:212m
C1R – module rKID:212m
C7 – module rKID:212m
module rKID:305
ZAP70 – module rKID:305
CD2 – module rKID:305
NKG7 – module rKID:305
CD247 – module rKID:305
module rKID:327
FGB – module rKID:327
FGG – module rKID:327
FGA – module rKID:327
TF – module rKID:327
Module EGS
Gene log2 FC
non–rejecting
MIXED
ABMR
TCMR
